# *Fus* depleted oligodendrocytes reduce neuronal damage and attenuate AD progression in the App^NL-G-F^ mouse

**DOI:** 10.1101/2025.11.24.689041

**Authors:** Te-Hsuan Tung, Sudhagar Babu, Xiangting Tang, Alec Sciutto, Micah Romer, Prathima Racha, Victoria Fisler, Adrita Saha, Takashi Saito, Takaomi C. Saido, Judith B. Grinspan, Hansruedi Mathys, Takashi D. Y. Kozai, Franca Cambi

**Affiliations:** Pittsburgh Institute for Neurodegenerative Diseases, School of Medicine, University of Pittsburgh, Pittsburgh, PA 15206, USA; Department of Neurobiology, School of Medicine, University of Pittsburgh, Pittsburgh, PA 15206, USA; Children Hospital of Philadelphia, Perelman School of Medicine at the University of Pennsylvania, Philadelphia, PA 19104, USA; Department of Neurocognitive Science, Institute of Brain Science, Nagoya City University, Nagoya, Aichi, Japan; Laboratory for Proteolytic Neuroscience, RIKEN Brain Science Institute, Wako-shi, Saitama, Japan; Department of Bioengineering, University of Pittsburgh, Pittsburgh, PA 15213, USA; Center for the Neural Basis of Cognition, University of Pittsburgh and Carnegie Mellon University, Pittsburgh, PA, United States of America; . Center for Neuroscience, University of Pittsburgh, Pittsburgh, PA, United States of America; . McGowan Institute for Regenerative Medicine, University of Pittsburgh, Pittsburgh, PA, United States of America; . Neuroscience Institute, Carnegie Mellon University, Pittsburgh PA, United States of America; Veterans Administration Pittsburgh, Pittsburgh, PA, 15213, USA

## Abstract

Oligodendrocytes (OL) and myelin abnormalities have emerged as important contributors to the pathogenesis of Alzheimer’s Disease (AD). OL maintain neuronal health through myelin axon interactions and by supplying neurotrophic and metabolic support. To gain insight on how OL and myelin may improve neuronal deficits associated with AD, we have generated a novel mouse model (AD/cKO) by crossing the App^NL-G-F^ mouse which carries three human AD mutations in the mouse *App* gene with the *Fus^OL^*cKO, which has thicker myelin associated with greater cholesterol biosynthesis. The spatial working memory deficits manifested by the aged AD mouse were fully rescued by the *Fus^OL^*cKO. This outcome was associated with reduced neuronal oxidative damage, preserved presynaptic structures at the plaque niches and a shift in microglia state at the niches in both hippocampus and cortex. In contrast, plaque burden and microglia density were decreased in the hippocampus but not in cortex, uncoupling the neuronal and microglia effects from the amyloid burden. Single cell transcriptomics of AD/cKO hippocampal OL revealed upregulation of energy metabolism and antioxidant genes, suggesting a role of OL enhanced energy metabolism in protecting neurons and affecting microglia state in AD pathology.

## Introduction

Oligodendrocytes (OL) the myelin producing cells of the central nervous system support neuron and neural network integrity and function by facilitating saltatory conduction through myelin deposition around axons, metabolic and trophic support to neurons^1^, maintaining axon integrity and function ^2–5^ and enhancing synapse formation ^6^. Loss of these critical functions supplied by OL and myelin has emerged as important contributor to the pathobiology of Alzheimer’s Disease (AD) ^7^.

AD is an age-dependent neurodegenerative disorder and represents the most common type of dementia in the aging population. Several lines of evidence converge to show that abnormalities in OL and myelin underlie AD pathogenesis. White matter abnormalities were identified by Magnetic Resonance Imaging in patients with Mild Cognitive Impairment amnestic and prodromal AD before the cognitive deficits manifest ^8–13^. Gene expression profiling studies in AD brain tissue have shown that OL are severely impacted early in the AD course and throughout disease progression ^14^ and suggest that disruption of OL integrity is a critical mechanism in AD pathogenesis and further corroborate the imaging and neuropathological findings.

Recent studies have investigated the role of OL and myelin in AD mouse models by either reducing myelin stability or enhancing new myelin formation^15,16^. Genetic and injury induced myelin loss worsened pathology in AD mouse models by interfering with microglia degradation of A β accumulations associated with a shift to an inflammatory microglia state without clear worsening of cognitive decline ^15^. By contrast, either genetically or pharmacologically induced oligodendrogenesis, reduced AD-related cognitive decline in an AD mouse model without correcting pre-existing myelin degradation, affecting plaque burden and microglia density ^16^. A selective vulnerability of myelin-axon interactions leading to disruption of myelin axon signalling was uncovered in both AD brains and mouse AD models^7^. These studies highlight a complex relationship between OL, myelin, neuronal function, amyloid deposition and communications with other glia cells in the context of AD pathology.

To resolve some of this complexity and advance our understanding of OL and myelin mediated neuroprotection, we have investigated whether OL equipped with greater myelin biosynthetic capacity can ameliorate AD pathology and rescue the cognitive deficits. We have leveraged the *Fus*^OL^cKO mouse that carries conditional depletion of Fused in Sarcoma (Fus) in OL ^17^. This mouse has greater myelin thickness and higher percentage of myelinated small caliber axons driven by enhanced cholesterol biosynthesis without an increase in OL cell number. Importantly, chronic electrophysiological recording *in vivo* in the *Fus*^OL^cKO visual cortex and the underlying hippocampus revealed a selective increase in visually evoked single-unit detectability and firing rate in the CA1 subregion relative to wild-type littermates^18^. The data indicates that *Fus* depleted OL support circuit integrity in the hippocampus a critical component of the memory loop.

In this study, we have generated a novel AD mouse model by crossing the App^NL-G-F^ mouse, a model of prodromal AD, with the *Fus*^OL^cKO and examined AD disease progression in the hippocampus and cortex of the App^NL-G-F^/ *Fus*^OL^cKO and App^NL-G-F^. *Fus*^OL^cKO rescued spatial working memory deficits manifested by the aged AD mice. This outcome was associated with reduced neuronal oxidative damage, preserved presynaptic structures and a shift in microglia state at the plaque niches in both hippocampus and cortex. In contrast, plaque burden and microglia density were decreased in the hippocampus but not in cortex, uncoupling the neuronal and microglia effects from the amyloid burden. *Fus* dependent myelin increase was present in both hippocampus and cortex. Single cell transcriptomics of AD/cKO hippocampal OL revealed upregulation of energy metabolism and antioxidant genes, suggesting a role of OL enhanced energy metabolism and myelin in protecting neurons and affecting microglia state in AD pathology.

## Results

### Spatial working memory is preserved in the aged AD/cKO and plaque burden is reduced in the hippocampus

The novel AD mouse model was generated by crossing the *App*^NL-G-F^, a prodromal model of AD^19^, with the *Fus*^fl/fl^/*Cnp*^cre/+^a developmentally and physiologically regulated hypermyelinating mouse model ^17^. The *App*^NL-G-F^ knockin mouse carries human AD mutations under the endogenous App promoter regulation and, unlike other mouse models, is not over-expressed and in the *Fus*^fl/fl^/*Cnp*^cre/+^, unlike other hypermyelinating mouse lines, the increase in myelin is developmentally regulated and additional myelin deposition does not continue unchecked past the completion of myelination ^17^. We reasoned that with these two genetic approaches, we could ask whether, in a physiological context, having more myelin and more robust OL can slow down or reverse AD disease progression and mitigate cognitive and pathological abnormalities.

We conducted all studies in the *App*^NL-G-F^ (referred to as AD), *App*^NL-G-F^/ *Fus*^fl/fl^/*Cnp*^cre/+^ (referred to as AD/cKO), *Fus*^fl/fl^/*Cnp*^cre/+^(referred to as cKO) and *Fus*^fl/fl^ (referred to as WT) with the WT and cKO serving as age matched controls and the cKO as genotype specific control for the AD/cKO (Fig. 1A). To examine the effects that *Fus*-depleted OL exert on the temporal and spatial progression of disease pathology, we have examined the hippocampus, critical component of the integrated memory circuit ^20–22^ and somatosensory cortex, which is involved in higher cortical functions, and selected the rostral S1 at the level of prefrontal cortex ^23^. A comprehensive battery of studies was performed at time points selected on the basis of previous characterization of Aβ deposition, microglia and astrocyte activation in the AD mouse ^19,24^ (Fig. 1A). At 2 months of age there is no A β accumulation, at 6 months plaques are formed, at 12- and 16-months full plaque load is present ^19,24^.

**Figure 1.**
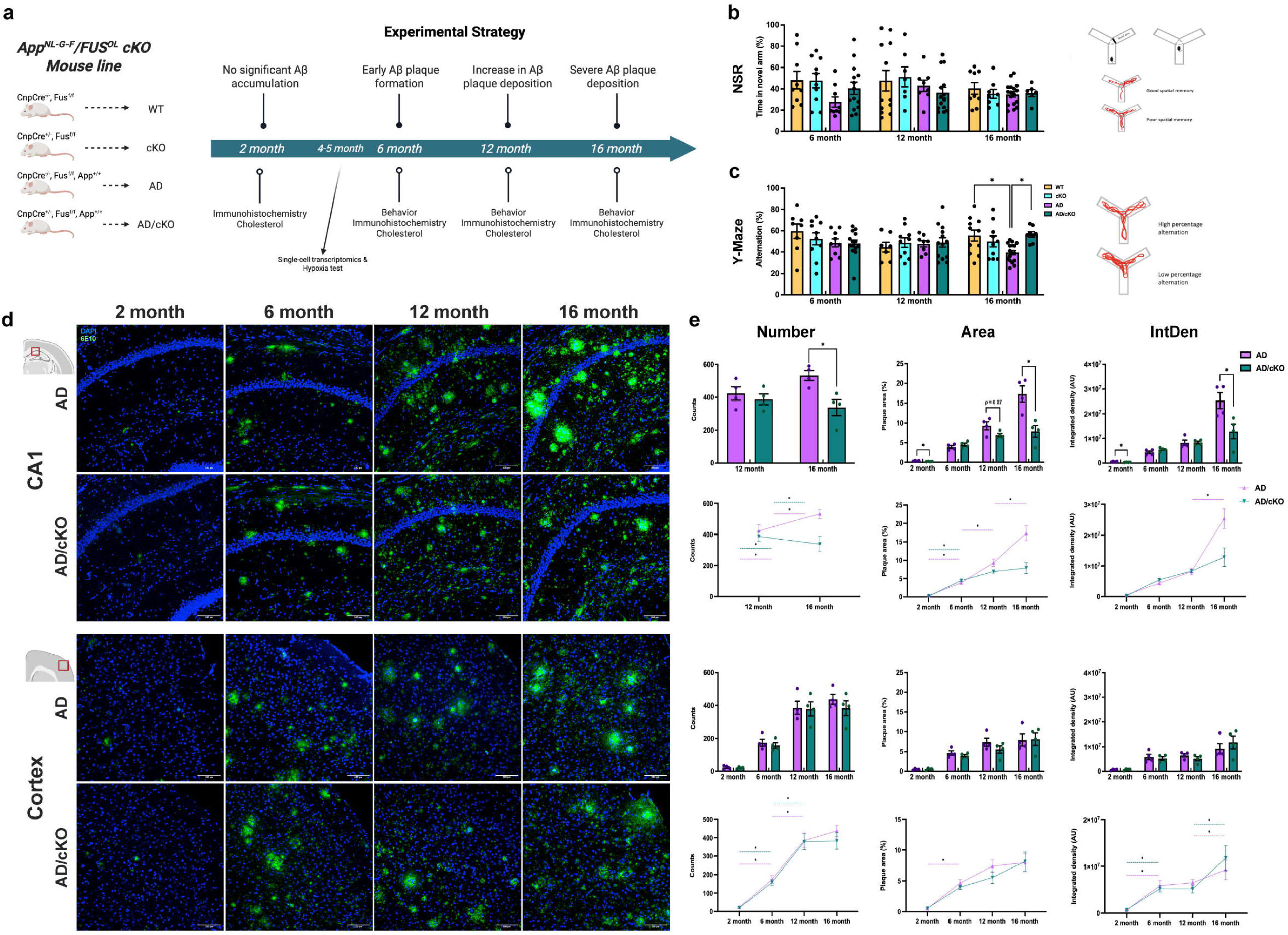
Aged AD/cKO mice preserve spatial working memory and show reduced plaque accumulation in the CA1 subregion of the hippocampus. **a.** Graphic representation of the mouse line indicates the four genotypes, WT, cKO, AD and AD/cKO and the Experimental Strategy shows the timeline and the experimental approaches. **b.** Bar graphs show the mean±SEM time (%) that each mouse spent in the novel arm after a trial period in which one of the arms was closed (Novel Spatial Recognition). There are no statistically significant differences between genotypes. The graphic representation shows the Y maze with the arm blocked used in trial 1 and the Y maze with the new arm open used in trial 2 and the read outs. **c.** Bar graphs show the the mean±SEM spontaneous alternations (%) in the Y maze for each mouse. The AD mice show a significant reduction in percent alternation vs. WT (39.47%±2.07 vs. 55.51%±5.16, p=0.0122). The percent alternation in AD/cKO mice shows no difference vs. WT and cKO (56.65%±2.54 vs. 55.51%±5.16, p=0.998; 56.65%±2.54 vs. 50.03%±5.20, p=0.697) and are significantly higher vs. the AD mice (56.65%±2.54 vs. 39.47%±2.07, p=0.016). The graphic representation shows the Y maze and the read outs. **d.** Representative 20X confocal images of cryosections stained with 6E10 to label 13 amyloid and DAPI to identify nuclei in the CA1 subregion and the sensory cortex of 2-, 6-, 12- and 16-months old AD and AD/cKO. Scale bar is 100µm. **e.** Bar graphs show the mean±SEM of the plaque counts, the area occupied by the plaques and the integrated density (intensity of fluorescence per area) per FOV. All three readouts are significantly lower in the CA1 of 16 months old AD/cKO vs. AD (counts: 338.2±48.4 vs. 532.0±29.9, p=0_014; area: 7.87%±1.48 vs. 17.33%±2.04, p=0.009; ln!Den: 1.28×107±3.01×106 vs. 2.53×107±3.19×106, p=0.029). No differences were detected at 6 and 12 months. No significant differences were detected in any of the readouts in the cortex of AD/cKO and AD at 6, 12 and 16 months. The rate of plaque accumulation was similar in the AD and AD/cKO up to 12 months in the CA1 and up to 16 months in the cortex.

Mice 6, 12 and 16 months old were tested for short-term spatial memory with the Novel Spatial Recognition (NSR) test by measuring the time mice spent in the novel arm following training in the Y maze with the arm blocked^25^. The time the AD and AD/cKO mice spent exploring the new arm was similar to WT and cKO at all ages tested (Fig. 1B), showing no decline in short term memory in either the AD or the AD/cKO. Spatial working memory was assessed with the spontaneous alternation test in the Y maze by measuring the number of entries into a new arm rather than re-entering the arm most recently visited (Fig. 1C) ^25^. At 16 months of age the AD mice showed statistically significant reduction in spontaneous alternations relative to age matched WT (Fig. 1C, p=0.012), while the AD/cKO performed at the same level as the WT (Fig. 1C) and cKO (Fig. 1C), indicating that spatial working memory is preserved in the AD/cKO vs. AD (Fig. 1C, p=0.016). WT and cKO performed similarly in all tests.

In previous studies, the AD mouse was shown to be hyperactive and to have reduced anxiety behavior^26^. We tested hyperactivity by counting the number of entries in the arms of the Y maze and the total distance travelled in the Elevated Plus Maze (EPM), and assessed anxiety in the EPM by measuring the time spent in the open vs. closed arm. The number of entries in the Y maze was significantly increased in both AD and AD/cKO relative to WT and cKO (SFig.1A, p=0.021). No statistically significant difference was detected between the AD/cKO and AD (SFig. 1A). Total distance travelled was also significantly higher in the AD and AD/cKO relative to WT and cKO (SFig. 1B, p=0.001). No statistically significant difference was detected between the AD and AD/cKO (SFig. 1B). The AD and AD/cKO spent more time in the open arm relative to WT and cKO (SFig. 1C AD vs. WT p=0.012; AD/cKO vs. cKO, p=0.0002), with the AD/cKO spending significantly more time than the AD (SFig. 1C, p=0.003). The data indicates that the *Fus*^OL^cKO enhances the anxiolytic effect of AD but not the hyperactivity, suggesting a selective effect on pathways underlying anxiety. Importantly, no differences were detected in either hyperactivity and anxiety between the WT and the cKO, as previously published^17^.

Next, we examined whether preserved working memory in the aged AD/cKO vs. AD was associated with reduced Aβ plaque burden, measured by number of plaques per field of view (FOV), area occupied by plaques and the integrated density, in the hippocampus and compared to cortex (Fig. 1D and E). No plaques were detected in 2 months old cortex and CA1 of the AD and AD/cKO (Fig. 1D and E). In CA1, plaque burden increased from 6 to 16 months (Fig. 1 D and E), but it was statistically lower in the AD/cKO compared to the AD at 16 months of age (Fig. 1 D and E, counts: p=0.014; area: p=0.009; IntDen: p=0.029). In the cortex, plaque accumulation increased progressively from 6 to 12 months and plateaued remaining stable at 16 months without statistically significant differences between AD and AD/cKO (Fig. 1 D and E). Next, we extended our analysis to the medial prefrontal cortex at the cingulate gyrus level (referred to as mPFC) which is implicated in an integrated circuit involved in spatial working memory^27^. The plaque number was not statistically different in the AD/cKO vs. AD mPFC (SFig. 4A), the plaque area and the integrated density were trending down in the AD/cKO (SFig. 4A). This data indicates that changes in plaque density is similar in different higher cortical function regions. Together, the data show that preserved spatial working memory at 16 months of age in the AD/cKO is associated with decreased plaque burden in the CA1, but not in cortical regions relevant to cognition.

### Microglia density and a shift to a protective phenotype are differentially regulated in the AD/cKO CA1 and cortex

Microglia activation is an early event in AD pathology ^28^ and microglia trajectories and states change in response to the environment and affect disease progression^29–32^. We have examined the temporal course of microglia cell accumulation by counting the number of Iba1+ cells in the cortex and CA1 of WT, cKO, AD and AD/cKO. In both AD and AD/cKO microglia density progressively increased in cortex and CA1 as the mice aged (Fig. 2 B, D). Iba1+ cell density was significantly higher in the CA1 of the AD vs. AD/cKO mice at 12 (Fig. 2A, B, p=0.0002) and 16 months of age (Fig. 2 A, B, p=0.0066). In contrast, no statistically significant differences were detected in cortex of the AD vs. AD/cKO at 12 and 16 months (Fig. 2 C, D). In the mPFC, the microglia density between the 16 months old AD/cKO and AD was not statistically significant (SFig. 4B).

**Figure 2.**
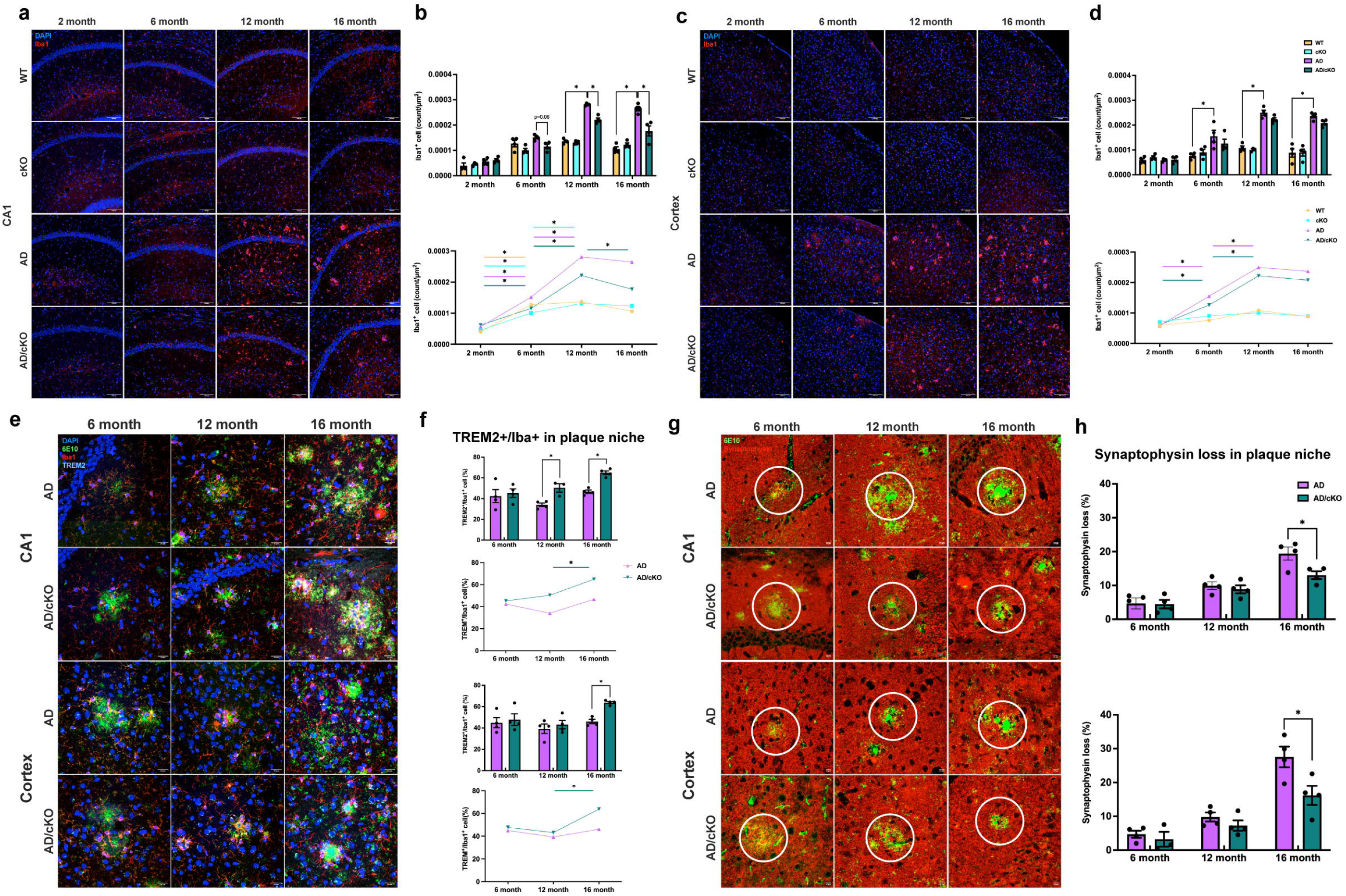
Analysis of microglia density, TREM2+ microglia at the plaque niche and pre-synaptic structures in CA1 and cortex. **a.** Representative 20X confocal images of lba1 stained cells in the CA1 of 2-, 6-, 12- and 16-months old WT, cKO, AD and AD/cKO. Scale bar is 100µm. **b.** Bar graphs represent the mean±SEM of lba1+ cell counts in the FOV of 3 sections in CA1 of 2-, 6-, 12- and 16-months old WT, cKO, AD and AD/cKO (n=4), *<0.05. Line plot shows the temporal course of lba1+ count increase in CA1 of 2-, 6-, 12- and 16-months old WT, cKO, AD and AD/cKO. **c.** Representative 20X confocal images of lba1 stained cells in cortex of 2-, 6-, 12- and 16-months old WT, cKO, AD and AD/cKO. Scale bar is 100µm. d. Bar graphs represent the mean±SEM of lba1+ cell counts in the FOV of 3 sections in cortex of 2-, 6-, 12- and 16-months old WT, cKO, AD and AD/cKO (n=4), *<0.05. Line plot shows the lba1+ count increase in the cortex of 2-, 6-, 12- and 16-months old WT, cKO, AD and AD/cKO. **e.** Representative 40X confocal images of 6E10 (green), lba1 (red) and TREM2 (cyan) stained CA1 and cortex of 6-, 12- and 16-months old AD and AD/cKO. White arrowheads point to TREM2+/lba1+ cells. Scale bar is 20µm. f. Bar graphs represent the mean±SEM of TREM2+/lba1 + at 3 plaque niches from 3 sections in CA1 and cortex (n=4), *p<0.05, **p<0.01. Line plot shows the temporal course of TREM2+/lba1+ percent increase at plaque niches of the CA1 and cortex of 6-, 12- and 16-months old AD and AD/cKO. g. Representative 40X confocal images of 6E10 (green), synaptophysin (red) stained CA1 and cortex of 6-, 12- and 16-months old AD and AD/cKO. Scale bar is 10µm. h. Bar graphs represent the mean±SEM of percentage loss of synaptophysin measured by lmaris rendering at 3 plaque niches from 3 sections in CA1 and cortex (n=4), *p<0.05.

To gain insight on the functional state of microglia, we have measured TREM2 (Triggering Receptor Expressed in Myeloid Cells) expression. TREM2+ microglia have stronger phagocytic capacity, cluster at the plaques where they are thought to reduce Aβ accumulations ^31,33^, in addition, they are anti-inflammatory and protective^30,34^. We measured TREM2+/Iba1+ microglia density at the plaque niche in 6-, 12- and 16-months old AD and AD/cKO CA1 and cortex. TREM2+/Iba1+ cells were statistically higher at the niche of the AD/cKO CA1 vs. AD at 12 (Fig. 2 E, F, p=0.009) and 16 months of age (Fig. 2 E, F, p=0.0003), while they were increased only at 16 months in the cortex (Fig. 2 E, F, p=0.0002) and mPFC (SFig. 4C p=0.0025) of the AD/cKO vs. AD. No statistically significant difference was present in 6 months AD/cKO vs. AD in cortex (Fig. 2E, F, p=0.70) and CA1 (Fig. 2E, F, p=0.72). Next, we asked whether the number of TREM2 microglia at the plaque niche in the AD/cKO is driven by higher total TREM2+ microglia and/or a shift from TREM2+ non plaque associated (NPAM) to plaque associated (PAM) microglia, defined as TREM2+ microglia associated with Aβ deposition. The total number of TREM2+, PAM and NPAM TREM2+ were not statistically different in 6-, 12- and 16-months old AD and AD/cKO CA1 (SFig. 2A) and cortex (SFig. 2B). TREM2+/Iba1+ microglia increased from 6 up to 12 months and plateaued (SFig. 2A, B). The data indicates that the higher number of TREM2+ microglia at the CA1 and cortex plaque niche of the AD/cKO is not accounted for by an increase in total TREM2+ microglia or an overall shift from the NPAM to PAM microglia. The data suggests that local signalling at the plaque niche affects TREM2+ microglia state ^35^.

Next, we sought to investigate how TREM2+ microglia may contribute to the improved memory function in the aged AD/cKO mice. While TREM2+ microglia limit Aβ depositions and may lead to preservation of memory by reducing plaque burden^31,36^, plaques were reduced only in CA1 but not in cortex despite the high number TREM2+ microglia at the niches. We postulated that TREM2 microglia exert a protective effect which is independent from plaque burden. To this end, we evaluated synaptic integrity by quantifying density of pre-synaptic structures in the AD/cKO vs. AD by synaptophysin staining, shown previously to be diminished at the plaques of the AD mouse ^19^. There is no statistically significant difference in synaptophysin loss at the plaque niche of the 6- and 12-months AD/cKO vs. AD CA1 and cortex (Fig. 2 G, H). In contrast, synaptophysin loss is significantly reduced at the plaque niche of 16 months old AD/cKO vs. AD CA1 (Fig. 2 G, H p=0.0295), cortex (Fig. 2 G, H, p=0.0333), and mPFC (SFig. 4 p=0.0008) suggesting that TREM2⁺ microglia are associated with reduced presynaptic loss independently from an effect on amyloid plaque burden.

### Astrocytic activation increases with disease progression in cortex and CA1 of the AD and AD/cKO

Astrocytes play a critical role in the maintenance of brain homeostasis by providing support to synapses especially in the hippocampus t^37–39^. Astrocytes undergo several morphological and gene expression changes in AD progression ^40^. We have characterized the temporal course of GFAP expression a marker of astrocyte activation associated with morphological cellular changes in the AD and AD/cKO. To capture GFAP+ cell bodies and processes we have measured the area occupied by GFAP+ staining in cortex and the CA1 (Fig. 3). Astrogliosis increases from 6 to 16 months in the AD (Fig. 3A, B, p<0.0001) and AD/cKO CA1 (Fig. 3A, B, p=0.0005) and in the cortex from 2 to 12 months when it plateaus in AD (Fig. 3 C, D, p<0.0001). In contrast, additional accumulation occurs between 12 and 16 months in the AD/cKO (Fig. 3 C, D, p<0.0001). The GFAP+ profiles were trending down in the CA1 of the 16 months AD/cKO vs. AD (Fig. 3A,B, p=0.35), while they were statistically higher in the cortex of the AD/cKO vs. AD (Fig. 3C, D, p=0.0002) and were trending up in the mPFC of the AD/cKO vs. AD (SFig.4D). An age-dependent increase in the GFAP+ profiles was present in both WT and cKO CA1 between 12 and 16 months (Fig. 3 A, B).

**Figure 3.**
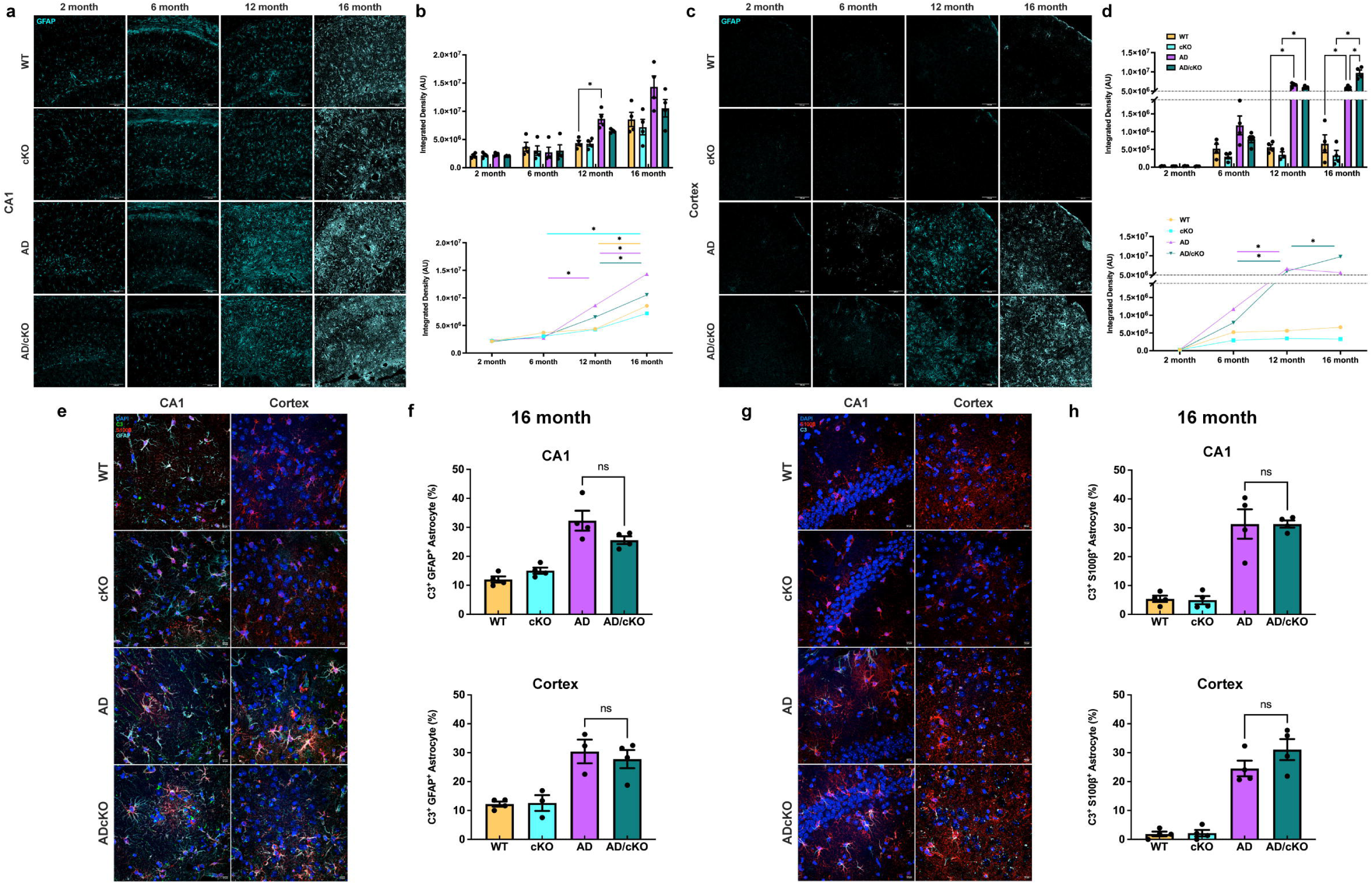
Analysis of astrocytic activation by GFAP expression and quantification of toxic astrocytes. **a.** Representative 20X confocal images of GFAP stained cells in the CA1 of 2-, 6-, 12- and 16-months old WT, cKO, AD and AD/cKO. Scale bar is 100µm. **b.** Bar graphs represent the mean±SEM of GFAP+ profile density (AU) in the FOV of 3 sections in CA1 of 2-, 6-, 12- and 16-months old WT, cKO, AD and AD/cKO (n=4), *<0.05. Line plot shows the temporal course of GFAP+ profile density (AU) increases in the CA1 *<0.05. **c.** Representative 20X confocal images of GFAP stained cells in the cortex of 2-, 6-, 12- and 16-months old WT, cKO, AD and AD/cKO. Scale bar is 100µm. **d.** Bar graphs represent the mean±SEM of GFAP+ profile density (AU) in the FOV of 3 sections in cortex of 2-, 6-, 12-and 16-months old WT, cKO, AD and AD/cKO (n=4), *<0.05. Line plot shows the temporal course of GFAP+ profile density (AU) increases in the cortex *<0.05. e. Representative 40X confocal images of GFAP (cyan) and C3(green) S10013 (red) stained CA1 and cortex of 16 months old AD and AD/cKO. Scale bar is 10µm. f. Bar graphs represent the mean±SEM of C3+/GFAP+ astrocytes in the FOV of 3 sections imaged at 20X in CA1 and cortex of 16 months old WT, cKO, AD and AD/cKO (n=4), ns= non-significant. **g.** Representative 40X confocal images of C3 (cyan) and S10013 (red) stained CA1 and cortex 16 months old AD and AD/cKO. Scale bar is 10µm. h. Bar graphs represent the mean±SEM of C3+/S10013+ astrocytes in the FOV of 3 sections imaged at 20X in CA1 and cortex of 16 months old WT, cKO, AD and AD/cKO (n=4), ns= non-significant.

GFAP+ astrocytes can be either homeostatic or cytotoxic ^41–43^. Next, we quantified the density of toxic astrocytes by measuring the expression of C3, a marker of toxic and inflammatory astrocytes ^41,42^ by staining with C3 and GFAP. We counted the number of C3+/GFAP+/DAPI+ cells in 16 months old AD/cKO and AD CA1 and cortex (Fig. 3 E, F). No difference was present in the percentage of C3+/GFAP+ astrocytes between AD and AD/cKO CA1 and cortex (Fig. 3 E, F).

Since fewer cortical astrocytes express GFAP compared to other brain regions, we stained with S100 β, a calcium-binding protein expressed broadly by cortical astrocytes ^44^. A statistically significant increase in C3+/S100β+ cells was detected in the CA1 of 16 months old AD relative to WT (Fig. 3G, H, p=0.0001), AD/cKO relative to cKO (Fig. 3 G,H, p=0.0001) and in cortex of AD vs. WT (Fig. 3G H, p=0.0001) and AD/cKO vs. cKO (Fig. 3 G, H, p<0.0001). The density of C3+/S100β + cells was not different in the AD/cKO vs. AD CA1 (Fig. 3G, H) and cortex (Fig. 3G, H). Taken together, the data shows no difference in toxic astrocyte density between AD and AD/cKO CA1 and cortex.

### Myelin density and cholesterol content are differentially regulated in the AD/cKO CA1 and cortex

The cKO has greater myelin thickness and cholesterol content in white matter tracts^17^, both supporting neuronal, axonal and synaptic health ^45–47^. We have measured compact myelin density by staining for MBP and PLP ^48^, and new myelin segment deposition by staining for MOG which resides in the outermost layer of myelin ^49^ ^50^. In the CA1, there was a physiological increase in MBP and MOG expression from 2 to 6 months in WT (Fig. 4 A,B,C, MBP, p=0.0004; MOG, p=0.0034), cKO (MBP, p=0.0003; MOG, p=0.0044), AD (MBP, p=0.0047; MOG, p=0.0003) and AD/cKO (MBP, p=0.0001; MOG, p<0.0001) with stable levels at 12 and 16 months of age (Fig. 4A, B, C). While the cKO and AD/cKO had greater MBP expression relative to WT and AD, consistent with *Fus* depletion driven increase in myelin thickness, the increase was statistically significant only in the cKO vs. WT at 6 (Fig.4 A, B, C, p=0.004) and 12 months (Fig. 4A B C, p=0.019). MOG was not increased consistent with its location in the outermost layer, hence, not affected by an increase in compact myelin thickness and indicating absence of new myelin segment deposition (Fig. 4 A, B, C, 2 month p=0.466; 6 month p=0.268; 12 month p=0.41; 16 month p=0.07).

**Figure 4.**
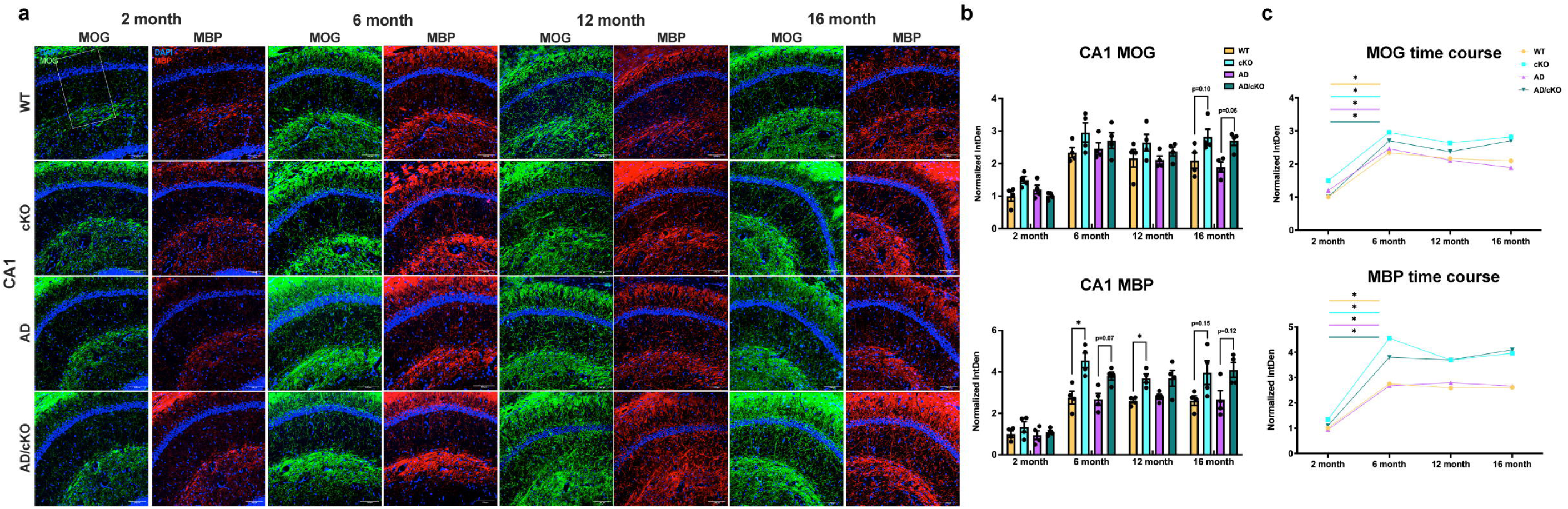
Fus depletion dependent increase in myelin profiles in AD/cKO hippocampal CA1. **a.** Representative 20X confocal images of MOG and MBP stained CA1 subregion of WT, cKO, AD, AD/cKO at 2, 6, 12 and 16 months. Scale bar is 100µm. **b.** Bar graphs represent the mean±SEM of the normalized density (MBP and MOG density at 2 months of age is set as 1) of MOG and MBP in the FOVof 3 sections in CA1 of 2-, 6-, 12- and 16-months old WT, cKO, AD, AD/cKO (n=4). *p<0.05, actual p value is shown for non-significant and trending down comparisons. c. Line plot shows the temporal course of normalized density of MOG and MBP at 2, 6, 12 and 16 months of age in WT, cKO, AD and AD/cKO.

In the L2-3 cortex, where a large proportion of axons are sparsely myelinated and adaptive myelination is easily detected in the adult brain^51,52^, MBP and MOG densities increased from 2 to 6 months in WT, cKO, AD and AD/cKO (Fig. 5 A,B,C). MBP and MOG densities were statistically higher in the cKO relative to WT (Fig. 5 A,B,C, MBP p=0.005; MOG, p= 0.003) and in the AD/cKO relative to AD (Fig. 5 A,B,C, MBP, p=0.0004 and MOG, p=0.014). A gradual decline in MOG occurred from 6 to 16 months, while MBP sharply declined at 12 moths (Fig. 5 A,B,C). In L4-6, where baseline axonal myelination is high, MBP and MOG densities were similar in all genotypes (SFig. 3A). The data suggests that *Fus* depletion in OL leads to increased myelin thickness and new myelin segment deposition in L2-3^49^.

**Figure 5.**
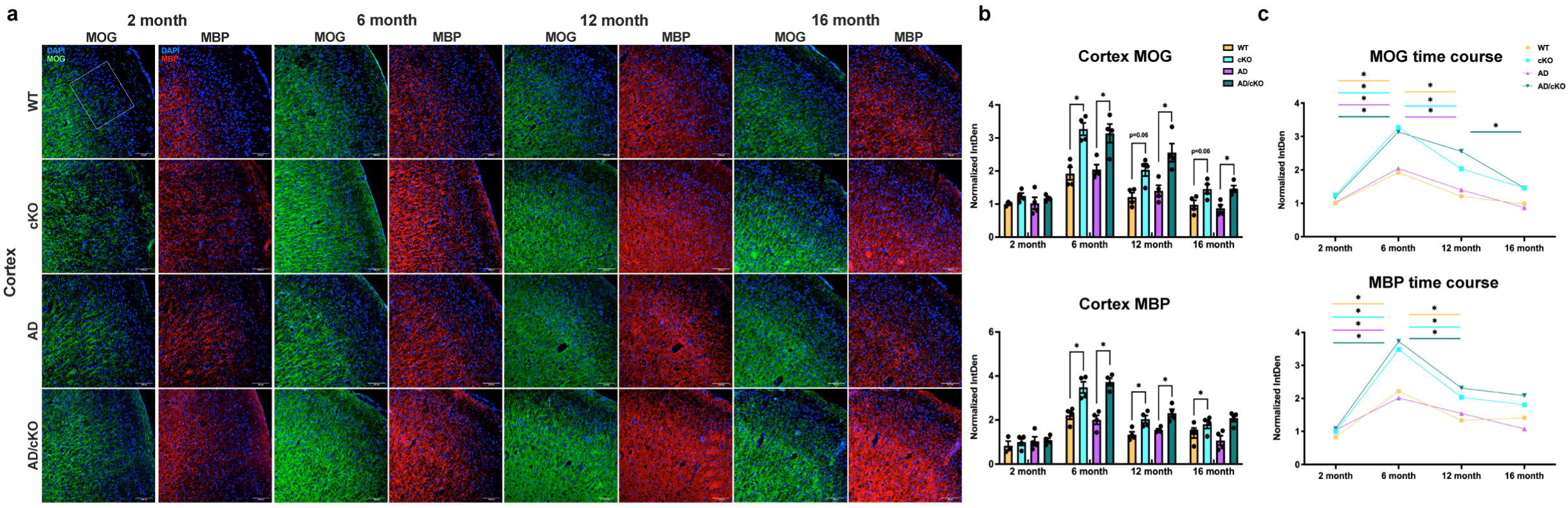
Fus-dependent increase in adaptive myelination profiles in the AD/cKO cortex. **a.** Representative 20X confocal images of MOG and MBP stained cortex L2-L3 of WT, cKO, AD, AD/cKO at 2, 6, 12 and 16 months of age. Scale bar is 100µm. **b.** Bar graphs represent the mean±SEM of the normalized density of MOG and MBP in the cortex L2-L3 ROI of 3 sections in 2, 6-, 12- and 16-months old WT, cKO, AD, AD/cKO (n=4). *p<0.05, ns= non-significant, actual p value is shown for trending down comparisons. **c.** Line plot shows the temporal course of normalized density of MOG and MBP at 2, 6, 12 and 16 months of age in WT, cKO, AD and AD/cKO.

Unlike MBP, the density of the major intrinsic myelin protein, PLP, in cortex L2-3 was not statistically different in the AD, AD/cKO vs. WT and cKO (SFig. 3B). Western blot analysis of PLP and MBP in cortex and hippocampus lysates of WT, cKO, AD and AD/cKO did not reveal any differences in protein levels (SFig. 3C, D). As previously shown in the cKO white matter ^17^, ASPA+ OL density was not increased in either cortex or CA1 of AD/cKO vs. AD and cKO vs. WT (data not shown).

Taken together, the studies did not reveal significant myelin loss in the AD vs. WT, in keeping with previously published data^15^, and in the AD/cKO vs. cKO. An increase in myelin measured by MBP immunostaining was present in CA1 and in the L2-3 of the AD/cKO and cKO cortex, while there was no statistically significant increase of PLP in L2-3 cortex.

Cholesterol content in higher in the cKO white matter tracts relative to WT mice ^17^.. Here, we have measured cholesterol and cholesterol esters with a colorimetric assay in lipids extracted from hippocampus and cortex of WT, cKO, AD and AD/cKO. Cholesterol was higher in the hippocampus of the cKO vs. WT at 2 months (Fig. 6, p=0.0095), 6 months (Fig. 6, p=0.0057) and 16 months (Fig. 6A, p=0.02) but not 12 month (Fig. 6A, p=0.133). Cholesterol content was trending up at 2 months AD/cKO vs. AD hippocampus (Fig. 6A, p=0.06) and was significantly higher in the AD/cKO vs. AD at 6 months (Fig. 6A, p<0.0001) but not at 12 and 16 months (Fig. 6A). No differences were present in the cortex of the four genotypes (Fig. 6B). The heterogeneity of MOL^53^ and regional restriction in lipid metabolism ^54^ may account for the spatial differences in cholesterol content between CA1 and cortex.

**Figure 6.**
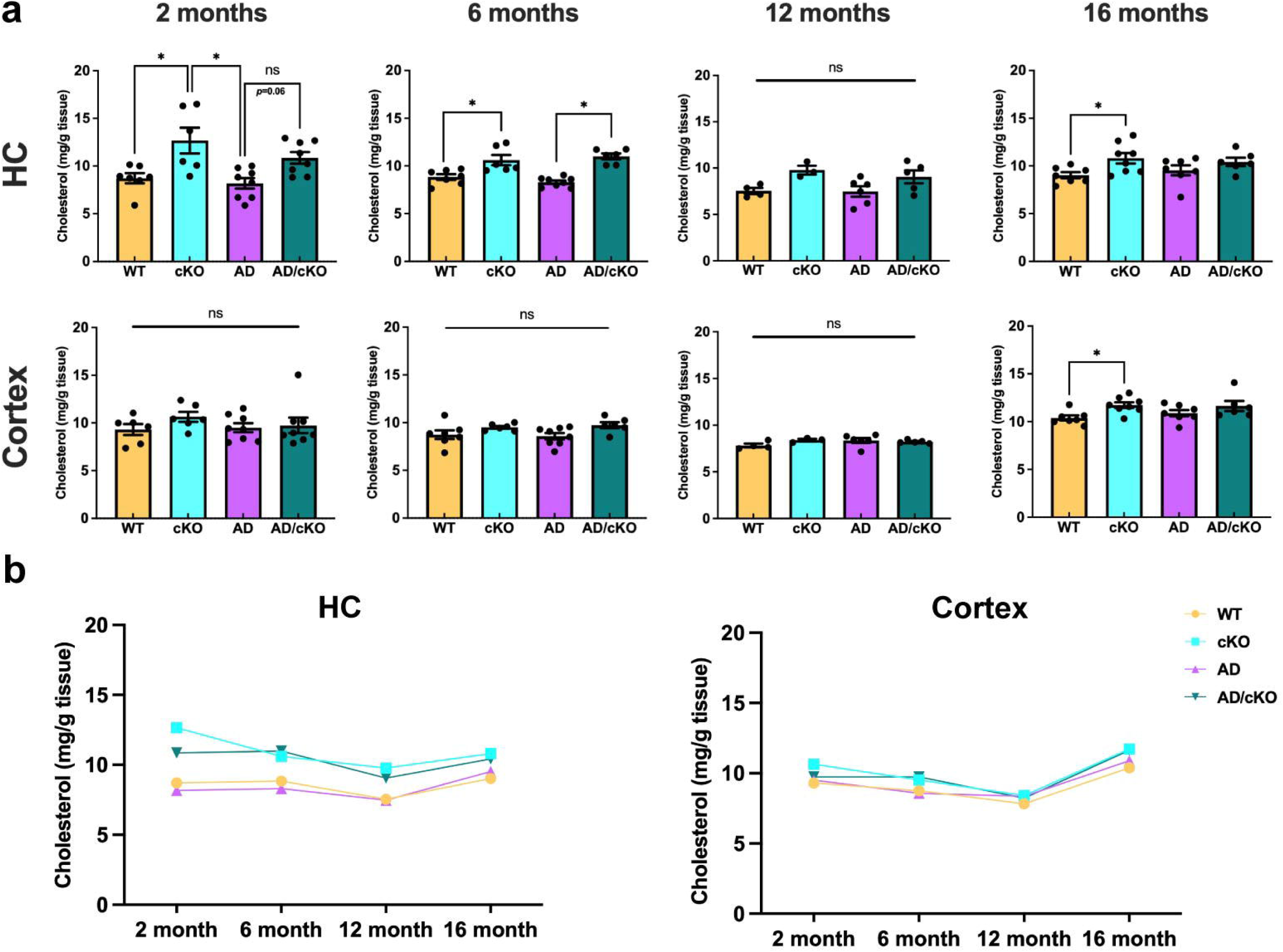
Cholesterol content is increased in the cKO and AD/cKO hippocampus. **a.** Bar graphs represent the mean±SEM of the cholesterol content (mg/g of wet tissue) from hippocampi and cortex collected from 2-, 6-, 12- and 16-months old WT, cKO, AD and AD/cKO brains, *p<0.05, ns= non-significant, actual p value is shown for trending up comparisons. **b.** Line plot shows the temporal course of the cholesterol content (mg/g of wet tissue) from hippocampi and cortex collected from 2-, 6-, 12- and 16-months old WT, cKO, AD and AD/cKO.

### Transcriptional profiles in AD/cKO hippocampus reveal significant gene changes in MOL and microglia

Our data has shown a broad impact of *Fus*-depleted OL on microglial cellular state, presynaptic structure loss at the plaque niche, and progression of AD pathology, particularly within the hippocampus, culminating in improved spatial working memory performance. To further elucidate how *Fus*-deficient OL mediate such widespread effects across diverse cell types, we performed a single-cell transcriptomic (SCT) analysis. Libraries were prepared from hippocampal cell suspensions isolated from WT, cKO, AD, and AD/cKO mice at 4–5 months of age (n = 4 per genotype; each sample pooled from two brains) using the 10X Genomics FLEX workflow (Fig. 7A). Following stringent quality control in accordance with current best practices, and subsequent clustering and annotation using reference datasets from the Allen Mouse Brain Atlas, we retained 115555 high-quality cells for downstream analysis. These cells were hierarchically classified into 26 major cell types, including telencephalic astrocytes (Astro-TE; 27,460 cells), microglia (32,980 cells), mature oligodendrocytes (MOL; 23,059 cells), and additional glial and hippocampal neuron-enriched populations such as DG Glut and DG-PIR Ex IMN (Fig. 7B, SFig.5A, B). Cell clusters were consistently represented across all samples, except for rare or less enriched populations such as non-telencephalic astrocytes (Astro-NT), arachnoid barrier cells (ABCs). As expected, we observed a marked depletion of *Fus* expression in MOL and a moderate reduction in *Fus* in committed oligodendrocyte precursors (COP), newly formed oligodendrocytes (NFOL), and myelin-forming oligodendrocytes (MFOL), in both cKO and AD/cKO compared to WT and AD mice (Fig. 7C).

**Figure 7.**
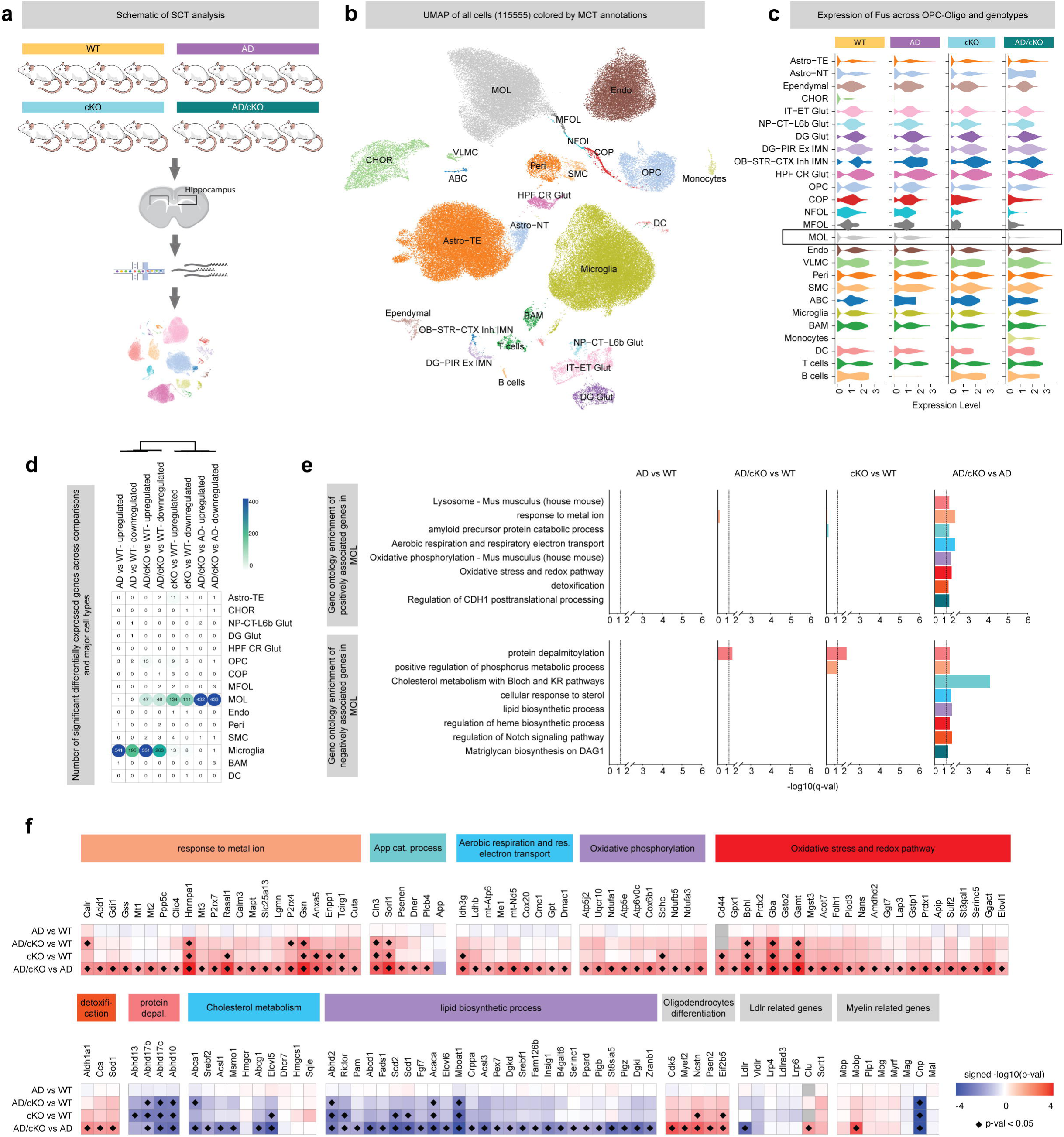
Mature oligodendrocytes exhibit pronounced transcriptomic alterations in AD/cKO compared with AD. **a**, Overview of the experimental design of SGT of hippocampi of WT, cKO, AD, and AD/cKO mice. **b**, UMAP visualization of the major cell populations identified by scRNA-seq. c, Violin plots showing Fus expression across major cell types and genotypes.d. Heatmap displaying the number of differentially expressed genes across genotype comparisons within each cell type. e. Gene ontology enrichment analysis for mature oligodendrocytes (MOL): the top panel highlights GO terms enriched among genes upregulated across genotype comparisons, while the bottom panel highlights GO terms enriched among downregulated genes. f. Heatmap showing the association of selected genes related to enriched Gene ontologies across genotype comparisons.

Differential gene expression analysis revealed that MOL and microglia exhibited the largest numbers of DEG. MOL were most affected in the *Fus* knockout contrasts (AD/cKO vs WT, cKO vs WT, AD/cKO vs AD), whereas microglial changes were evident in AD and AD/cKO comparisons versus WT (Fig. 7D). Specifically, in AD/cKO MOL compared to AD MOL, we identified 432 upregulated and 433 downregulated genes. In the cKO vs. WT comparison, 134 genes were upregulated and 111 downregulated in MOL. In contrast, only 47 genes were upregulated and 48 downregulated in AD/cKO MOL relative to WT (Fig. 7D). These findings indicate a strong impact of *Fus* depletion specifically in the AD background, suggesting a genotype-to-genotype interaction. Gene Ontology (GO) enrichment analysis of the DEG in AD/cKO vs. AD MOL revealed significant upregulation of genes involved in metal ion response, amyloid precursor protein catabolic processes, oxidative phosphorylation, aerobic respiration, respiratory electron transport, redox homeostasis, and detoxification pathways (Fig. 7 E, F). Conversely, genes associated with cholesterol metabolism and lipid biosynthesis were significantly downregulated. Additionally, genes related to *protein depalmitoylation* were downregulated in cKO vs. WT MOL (Fig. 7 E, F).

In microglia, we identified 541 upregulated and 196 downregulated genes in AD mice relative to WT, and 561 upregulated and 263 downregulated genes in AD/cKO mice relative to WT (Fig. 7D). GO enrichment analysis showed that most upregulated genes in both AD vs. WT and AD/cKO vs. WT comparisons were associated with inflammatory responses and cellular activation pathways, indicating a robust App-dependent transcriptional program (SFig. 5C, D).–Downregulated genes were enriched for pathways related to the regulation of protein phosphorylation Most of the up and downregulated genes in MOL are unique to the AD/cKO MOL vs. cKO MOL. Remarkably, oxidative phosphorylation genes of mitochondrial complex 1 (Ndufa1, 3 and Ndufb5, mt-Nd5), antioxidant (SOD1, Gstp1, Gss), electron transport chain are exclusively upregulated in the AD/cKO vs. AD but not in the cKO vs. WT (Fig. 7F). Some of the genes involved in OL differentiation such as Ncstn and Anxa5 are upregulated in both AD/cKO and cKO MOL (Fig. 7F), whereas others such as Myef2, Cdk5 and Psenen are exclusively up regulated in AD/cKO MOL, suggesting a stronger maintenance of OL identity and differentiation vs. the AD MOL. Interestingly, Ncstn and Psen2 that regulate processing of proteins, such as Notch important for OL differentiation are also involved in App processing, suggesting a potential cross regulation of App and other membrane protein processing ^55,56^. Finally, genes such as gelsolin which participates in myelin wrapping ^57^, and Cln3 involved in membrane, lysosome trafficking and cytoskeleton organization are up regulated in AD/cKO vs. AD and vs. WT and cKO vs. WT (Fig. 7F).

The SCT have revealed a significant down regulation of genes involved in fatty acid (FA) metabolism, fatty acid chain elongation and desaturation and sterol export. Interestingly, Elovl5, Elongase of PUFAs C3 and C6 polyunsaturated, Scd1 and Scd2, desaturases involved in synthesis of monounsaturated FAs and Acaca enzyme involved in rate limiting step in FA biosynthesis are downregulated in both AD/cKO and cKO MOL. Some of these genes fall into the GO enrichment category of cholesterol metabolism by affecting FA synthesis predominantly, however, the main cholesterol synthesis enzymes, namely Hmgcr and Hmgcs are not downregulated (Fig. 7F). The significance of these changes remains to be investigated, however, changes in FA saturation and side chain length will affect phospholipid and glycolipid side chains and may lead to changes in membrane fluidity ^58^.

To further determine whether these gene expression changes reflected altered microglial and MOL subtype proportions, we resolved the dataset into 37 high-resolution cell types, capturing seven MOL subclusters (MOL_1 to MOL_7) and six microglial subclusters (Microglia_1 to Microglia_6) (SFig. 6A, B). Microglia_1 through Microglia_3 expressed high levels of homeostatic markers such as *P2ry12* and *P2ry13*, whereas Microglia_4 and Microglia_5 exhibited reduced expression of homeostatic markers and increased expression of activation-associated genes, including *Lpl* and *Itgax*, indicative of activated states. Microglia_6 displayed a proliferative signature, marked by elevated *Mki67* expression. Clusters Microglia_4 and Microglia_5 were enriched in both AD and AD/cKO groups (SFig. 6B), identifying them as disease-associated microglial states (SFig. 6C). Proportional analyses revealed a significant increase in Microglia_4 and Microglia_5 in both AD and AD/cKO relative to WT, accompanied by a significant reduction in the homeostatic Microglia_1 population (SFig. 6D), with no significant differences between AD and AD/cKO (SFig. 6D). The seven MOL clusters exhibited distinct transcriptional signatures (SFig. 6C). MOL_3 showed elevated expression of genes involved in oligodendrocyte differentiation (*Enpp6*) ^59^ and cytoskeletal organization (*Anln*). MOL_7 was characterized by high expression of *Cntn1* (contactin), which is essential for myelin–axon interactions, as well as *Man1a* and *Pcdh7*, markers associated with late-stage, remyelinating mature oligodendrocytes (bioRxiv, Dolan et al., 2025). MOL_6, defined by expression of *Apoe* and metallothionein genes (*Mt2*, *Mt3*), was significantly increased in cKO mice relative to WT (SFig. 6C, D). Yet, MOL cluster proportions did not differ significantly between AD and AD/cKO groups (SFig. 6D). Taken together, the SCT data have revealed a major role of DEG in AD/cKO vs. AD in energy metabolism, antioxidant, App catabolism and OL differentiation and have uncovered a potential role of these genes in reducing neuronal damage and improving the homeostatic environment.

### Oxidative damage is reduced in the AD/cKO compared to AD

The upregulation of mitochondrial complex 1 and antioxidant genes (Fig. 7D) indicated a greater bioenergetics strength and resilience to oxidative stress of the AD/cKO MOL vs. AD. We reasoned that these gene changes may increase the ability of MOL to support neuron health and reduce oxidative damage. To test this hypothesis, we have examined whether hypoxic damage caused by Aβ-oligomer driven capillary vasoconstriction in the early stages of AD is reduced in the AD/cKO ^60,61^ by in vivo injection of pimonidazole (hypoxyprobe) followed by detection of pimonidazole conjugates with SH protein groups in neurons. The number of hypoxic neurons was increased in both AD and AD/cKO vs. WT and cKO at 4 months of age (Fig. 8A). While the count of hypoxic neurons was trending down in CA1 (Fig. 8A, p=0.17) and cortex (Fig. 8A, p=0.21) of the AD/cKO vs. AD it did not reach statistical significance.

**Figure 8.**
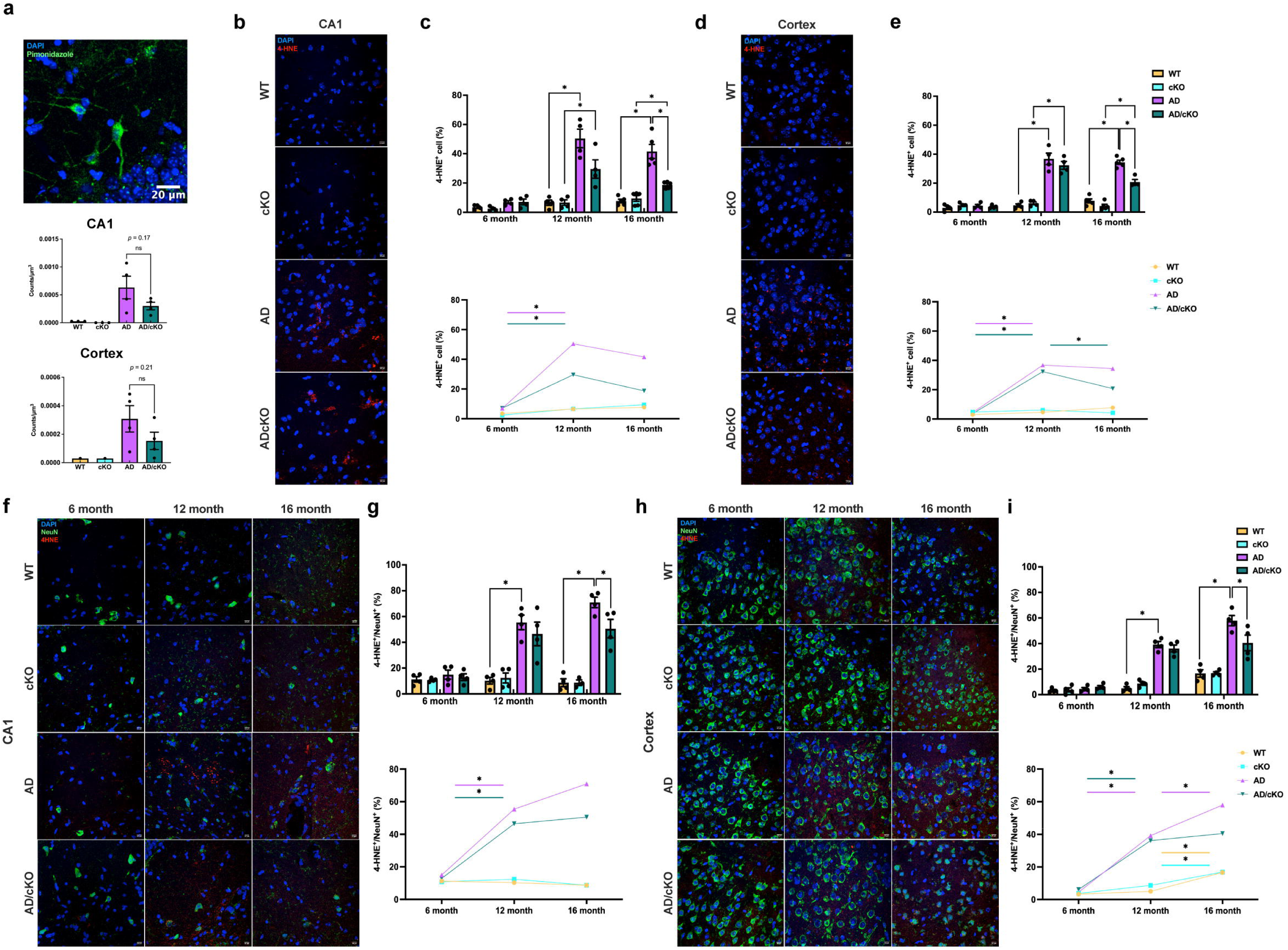
Oxidative damage measured by lipid peroxidation is reduced in the CA1 and cortex of the aged AD/cKO. **a.** Representative 20X confocal image of hippocampus neurons stained with an Ab to pimonidazole SH adducts of 4 months old AD. Bar graphs represent the mean±SEM of hypoxic neurons/µm3 in the CA1 and cortex in the FOV of 3 sections of WT, cKO (n=2}, AD and AD/cKO (n=4), actual p value is shown for non-significant and trending down comparisons. Scale bar is 20µm. **b.** Representative 40X confocal images of 16-months old WT, cKO, AD and AD/cKO CA1 stained with DAPI and an Ab for 4-HNE. Scale bar is 10µm. **c.** Bar graphs represent the mean±SEM of the 4-HNE+/DAPI+ cells(%) in the FOV of 3 sections imaged at 20X (n=4}, *p<0.05. Line plot shows the temporal course of the percentage of 4-HNE+/DAPI+ cells at 6, 12 and 16 months of age in WT, cKO, AD and AD/cKO. d. Representative 40X confocal images of 16-months old WT, cKO, AD and AD/cKO cortex stained with DAPI and an Ab for 4-HNE. Scale bar is 10µm. **e.** Bar graphs represent the mean±SEM of the 4-HNE+/DAPI+ cells(%) in the FOV of 3 sections imaged at 20X (n=4}, *p<0.05. Line plot shows the temporal course of the 4-HNE+/DAPI+ cell density at 6, 12 and 16 months of age in WT, cKO, AD and AD/cKO. f. Representative 40X confocal images of 6, 12 and 16 months old WT, cKO, AD and AD/cKO CA1 stained with an Ab to NeuN and an Ab to 4-HNE. Scale bar is 10µm. **g.** Bar graphs represent the mean±SEM of the 4-HNE+/NeuN+ cells(%) in the FOV of 3 sections imaged at 20X (n=4}, *p<0.05. Line plot shows the temporal course of the 4-HNE+/NeuN+ cells at 6, 12 and 16 months of age in WT, cKO, AD and AD/cKO. h. Representative 40X confocal images of 6, 12 and 16 months old WT, cKO, AD and AD/cKO cortex stained with Abs to NeuN and 4-HNE. Scale bar is 10µm. i. Bar graphs represent the mean±SEM of the 4-HNE+/NeuN+ cells in the FOV of 3 sections imaged at 20X (n=4), *p<0.05. Line plot shows the temporal course of the 4-HNE+/NeuN+ cells at 6, 12 and 16 months of age in WT, cKO, AD and AD/cKO.

To assess reactive oxygen species (ROS) damage, we measured lipid peroxidation by quantifying 4-HNE+ adducts^62–64^ in 6, 12, 16 month old WT, cKO, AD and AD/cKO. No increase of 4-HNE+ cells identified by DAPI was detected in the 6 months old AD and AD/cKO CA1 (Fig. 8 C) and cortex (Fig. 8 E) vs. WT and cKO. The 4-HNE+ cells were significantly increased in 12 months and 16 months CA1 and cortex of AD vs. WT and of AD/ckO vs. cKO (Fig. 8 C, E). The 4-HNE+ cells were statistically lower in the 16 months old AD/cKO vs. AD CA1 (Fig. 8 B,C, p=0.0001) and cortex (Fig. 8 D,E, p<0.0001). 4-HNE+ cells were very low in WT and cKO. The data indicate reduced oxidative damage in the AD/cKO.

Next, we assessed neuron oxidative damage in CA1 and cortex of 6-, 12- and 16-months old WT, cKO, AD and AD/cKO. No increase in the 4-HNE+/NeuN+ was detected in the 6 months old AD and AD/cKO CA1 (Fig. 8F, G) and cortex (Fig. 8H, I) vs. WT and cKO. At 12 and 16 months of age, the 4-HNE+/NeuN+ cells were significantly increased in CA1 and cortex of the AD vs. WT and AD/cKO vs. cKO (Fig. 8 G, I). The 4-HNE+/NeuN+ cells were significantly higher in the AD cortex 16 months old vs. 12 months old (Fig. 8H, p=0.0025), while no increase was observed in the AD/cKO (Fig. 8 H, I). A statistically significant reduction in 4-HNE+/NeuN+ cells was detected in the 16 months old AD/cKO CA1 (Fig. 8F, G p=0.021) and cortex (Fig. 8H, I p=0.04) vs. AD. Taken together, the data indicate that oxidative damage is decreased in neurons of the AD/cKO, suggesting a protective effect of *Fus* depleted OL.

## Discussion

Disruption of oligodendrocyte and myelin support to neurons are important contributors to AD pathobiology and represent potential targets for disease modification strategies. The current studies in a novel AD mouse carrying *Fus* depletion in OL have uncovered enhanced energy metabolism in MOL which is associated with restoration of spatial working memory. The robust functional memory improvement in the AD/cKO is evidence of a neuroprotective effect of Fus depleted OL. The SCT studies have revealed an unexpected upregulation of energy metabolism and stress related genes in *Fus* depleted AD MOL leading to a new understanding of how OL may ameliorate AD neurodegeneration and shifting the focus on the energy and metabolic support of MOL to neurons and other cells such as microglia and vessels. The upregulation of energy metabolism and antioxidant genes is present exclusively in AD/cKO MOL and not in the parent *Fus*cKO MOL, suggesting a genotype-to-genotype interaction and occurs early in disease prior to plaque accumulation, suggesting an adaptive response to enhance energy supply and antioxidant defence mechanisms. In a previous study, an Olig specific gene module characterized by up regulation of myelin genes and OL regulatory genes^24^ was identified by spatial transcriptomics of the App^NL-G-F^ at early stages of disease, representing an attempt of OL to support neurons with more myelin deposition. However, in advanced stages of disease, this response was lost indicating a failure of the OL to sustain cell function. The upregulation of energy metabolism and antioxidant genes revealed by our SCT studies suggest that AD/cKO MOL may be better equipped to sustain trophic support to neurons through advanced stages of disease. In keeping with this affirmation, rescue of neuronal function, clustering of more protective microglia and synapse preservation at plaque niches occur in the aged AD/cKO.

While we did not identify changes in myelin gene expression by SCT, these studies revealed upregulation of genes involved in OL differentiation and membrane transport that play a pivotal role in the maintenance of the OL differentiation state and the transport of myelin proteins to the myelin membranes. A *Fus* dependent increase in compact myelin was identified in both AD/cKO hippocampus and cortex L2-3 by MBP quantification and MOG increase in cortex L2-3 consistent with adaptive myelination in these cortical layers where a high number of axons are unmyelinated and partially myelinated relative to L4-L6 ^52^. Unlike MBP, PLP quantification did not reveal differences between the AD/cKO vs. AD and cKO vs. WT cortex L2-3. Since PLP is the most abundant and stable myelin protein, detection of changes may require a large effect size and/or may need a more quantitatively sensitive method of analysis.

The impact of Fus depleted MOL is broad and affects several cellular responses and disease pathology. Clustering of TREM2+ microglia at the aged plaque niches ^35^, was increased in the AD/cKO CA1, mPFC and cortex and was associated with preservation of presynaptic structures in all these regions. TREM2 microglia limit amyloid deposition ^31^, however, a reduction of plaque burden is present only in the AD/cKO hippocampus but not in cortex and mPFC, uncoupling plaque burden from synaptic structure preservation and rather suggests an effect of TREM2+ secreted anti-inflammatory cytokines^34,65,66^ and TREM2 mediated neuroprotection ^67^. The lower plaque burden in the hippocampus may result from the earlier TREM2+ microglia clustering and phagocytosis or reflect a different rate of Aβ production and deposition ^68^. OL account for 30% of Aβ amyloid production^68^, however, MOL subclusters are differentially represented in different brain regions^53^ and may have different Aβ production capacity. Interestingly, hippocampal *Fus* depleted AD MOL have reduced App gene expression (Fig. 7, p=0.00509) and may secrete less Aβ.

What drives the greater TREM2+ clustering at the plaque niche remains to be elucidated. Since there is no shift in non-plaque associated to plaque associated TREM2+ microglia ^69^, we postulate that local signals at the plaque niche regulate TREM2+ cell biology by enhancing the maintenance of the activated state and preventing the microglia acquisition of a senescent state ^70^. The niche microenvironment fosters signalling via ligand receptor pair expression ^35^. A major signalling ligand-receptor pair is the Csf1-Csf1r via autocrine mechanism ^35^ and heterologous signalling from astrocytes and OL ^71,72^. Since OL are a major source of Csf1 in adult brain and development ^71^, it is possible that *Fus* depleted MOL at the plaque niche secrete Csf1 and promote TREM2+ microglia ^70^. The SCT studies performed prior to plaque deposition, revealed upregulation of Csfr1 and Csf1 in both AD and AD/cKO microglia indicating an activation of microglia, but there was no upregulation of Csf1 in MOL (data not shown). However, MOL at the plaques may have different gene expression profiles that are not captured by single cell transcriptomics especially prior to plaque deposition. Future spatial transcriptomics in the aged mice will elucidate MOL to microglia signalling in the AD/cKO plaque niche.

Increased reactive oxygen species production is associated with aging and AD ^73^. The sharp increase of neuronal oxidative damage at 12 and 16 months is likely to result from neuron intrinsic energy, antioxidant failure and loss of extrinsic support from glial cells. The amelioration of neuronal oxidative damage in the aged AD/cKO suggests that the energetically strong MOL support neuron and axon viability. Hypoxia caused by pericyte mediated capillary constriction is an early event in AD, which leads to hypoxic damage to neurons ^60,61,74^. Mature oligodendrocytes associate directly with blood vessels in both cortex and hippocampus, are an integral component of and contribute to the integrity of the blood vessel wall ^75–77^. A downward trend in the number of hypoxic neurons in the AD/cKO hippocampus and cortex, although not statistically significant, suggests a potential beneficial effect of AD/cKO MOL in reducing hypoxia possibly by improving vascular health.

Lipids are critical for myelin maintenance and neuron and synapse health ^78^. The studies have revealed that cholesterol content is higher in the hippocampus of the cKO up to 16 months and up to 6 weeks in the AD/cKO. In contrast, the cholesterol content is not increased in the cortex of cKO and AD/cKO. The spatial difference between cortex and hippocampus, a mixed white/grey matter region, may reflect regionally regulated myelin cholesterol and lipid composition ^54^.

The SCT studies have revealed down regulated genes involved in Fatty Acid biosynthesis, side chain elongation and protein depalmitoylation and have uncovered a previously unknown role of these lipid pathways and protein modifications. The changes in the FA side chains of phospholipids and glycolipids are predicted to change membrane fluidity and affect membrane protein transport^58^. Changes in palmitoylation are likely to affect protein interactions ^79^. Future studies will elucidate the impact of these changes on AD pathology and progression.

In summary, we show that higher myelin density induced by *Fus* depletion and upregulation of energy metabolism and antioxidant gene expression induced by *Fus* depletion in the context of App gene mutations are associated with reduced oxidative damage in neurons and a more homeostatic environment by affecting microglia state at the plaque niche.

## Materials and Methods

### Animals and Experiment Design

#### Ethics Statement

The use of animals in this study followed the guidelines of the Institutional Animal Care and Use Committees (IACUC) of the University of Pittsburgh (Animal Welfare Assurance number D16-00118) and was approved by IACUC under protocol # 24034725.

#### Generation of the App^NL-G-F^/Fus^OL^cKO line and Genotyping

The homozygous *App^NL-G-F^* mouse line (a kind gift of Dr. Saido^19^) was crossed with the *Fus*^fl/fl/CNPcre/+^ line ^17^ to generate the *App^NL-G-F^*/*Fus*^fl/fl/CNPcre/+^. This new line carries App*^NL-G-F^* and FUS^fl/fl^ in homozygosity and CNP^cre/+^in heterozygosity yielding *App^NL-G-F^/Fus*^fl/fl/CNPcre/+^(*App^NL-G-F^/Fus^OL^cKO)* and *App^NL-G-F^/Fus*^fl/fl/CNP+/+^(App*^NL-G-F^*). The new line was maintained by breeding *App^NL-G-F^/Fus*^fl/fl/CNPcre/+^ with the littermates *App^NL-G-F^/Fus*^fl/fl/CNP+/+^. For simplicity, we will refer in the text to *App^NL-G-F^/Fus^OL^cKO* as AD/cKO and *App^NL-G-F^/Fus*^fl/fl/CNP+/+^ as AD. The original *Fus*^OL^cKO line was maintained by breeding *Fus*^fl/fl^/^Cnpcre/+^ with the *Fus*^fl/fl^ littermates ^17^. We will refer to *Fus*^OL^cKO as cKO and *Fus*^fl/fl^ as WT. The four genotypes were used for the experiments described in this paper (Fig. 1A). All mice were bred and housed in a barrier facility with a 12-hour light/dark cycle. Food and water were available ad libitum. Mouse genotypes were determined by PCR of ear snip-derived DNA following previously published protocols for *Fus*^fl/fl^ and Cnp^cre^ ^17^ and *App^NL-G-F^*^19^. Both males and females were used for all the analyses. All efforts were ^22^made to minimize animal suffering and the number of animals used.

#### Mouse Behavioral testing

All behavioral data were collected in the Preclinical Phenotyping Core of the University of Pittsburgh School of Medicine.

#### Novel Spatial Recognition

To assess short-term recognition memory, we used the novel spatial recognition test which is similar to the novel object recognition but replaces the use of objects with visual cues ^25,80^. Mice were acclimated to the room under ambient lighting conditions (50 lux) at least 1 hour prior to actual test. White noise generators were used to minimize background noise in the testing room (∼60-70dB). Mice were placed within a three-arm Y-shaped maze (27 cm length, 6 cm width, and 15 cm heigh, clear polycarbonate), in which one of the arms was inaccessible, surrounded by a black floor-to-ceiling curtain to minimize visual cues outside the maze. The three arms are defined as start arm, familiar arm, and novel arm. The start arm is at the entrance of the perimeter curtain. A solid black polycarbonate wall (6 cm width, 15 cm height) is placed at the entrance to the novel arm to prevent access to that arm during the first trial. Mice were allowed to freely explore for 10 minutes and then returned to their home cage. Mice are given a delay period of 10 min prior to being placed back in the three-arm maze with the barricade removed, giving access to all three arms. Mice were allowed to freely explore for 5 minutes. Behavior was recorded and the amount of time spent in each arm and the number of entries into each arm were tracked (EthoVision XT 9.0, Noldus Information Technology, Wageningen, Netherlands), exported and used for the analysis. To calculate the total arm entries per mouse, we summed the number of entries into each arm. From the cumulative time spent in each arm, the percent duration in each arm was calculated 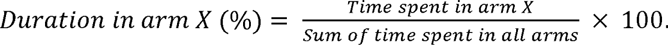 The preference in spending time in the novel arm relative to the familiar or start arms was taken as a measure of short-term memory.

### Spontaneous Alternation

To assess spatial working memory and exploration, we tested mice in the Y Maze ^81–83^. Mice were acclimated to the room under ambient lighting conditions (50 lux) at least 1 hour prior to actual test. White noise generators were used to minimize background noise in the testing room (∼60-70dB). A Y-shaped maze (33.65 cm length, 6 cm width, 15 cm height, clear polycarbonate) placed on top of an infrared backlit background (Noldus Information Technology, Wageningen, Netherlands) surrounded by a black floor-to-ceiling curtain to minimize visual cues outside the maze. When placed within a maze with multiple arms, mice exhibit the spontaneous behavior of alternating between arm choices with a greater frequency than re-entering the same arm most recently visited ^83^.The mouse was placed at the end of the start arms of the maze facing the maze center and allowed to explore freely for 8 min. Behavioral tracking software (EthoVision XT 9.0, Noldus Information Technology, Wageningen, Netherlands) was used to automatically track the behavior of the center point of the mouse’s body to determine the order of arm entries. From the sequence of arm entries, the number of alternation opportunities were counted as the total number of correct and incorrect triads. The percent alternation was calculated as the ratio of correct triads to the number of alternation opportunities: 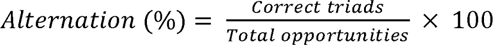. The correct alternations when the mouse entered three different arms consecutively was taken as a measure of spatial working memory ^25^.

### Elevated Plus Maze (EPM) test

To assess anxiety behavior we used the elevated plus maze ^84^. Mice were acclimated for 30 minutes prior to testing in a well-lit room. The mouse was placed in a closed arm of the EPM) which has two closed arms and two open arms and was allowed to explore the apparatus for 5 min. Experiments were conducted at full light. All movements were recorded by an overhead camera using SMART 3.0, Panlab S.I. All the videos were collected and automatically analysed by behaviour tracking system (SMART 3.0, Panlab S.I., Barcelona, Spain). We measured time spent in the open arms as a measure of anxiety and the total distance traveled, as a measure of activity. Time spent in the open arms was taken as an inverse measure of anxiety.

#### Lipid extraction and Cholesterol quantification

Mice were deeply anesthetized, perfused with cold PBS and rapidly decapitated. Brains were removed and prefrontal cortices and hippocampi were carefully dissected out on ice ^85^. All tissues were rapidly frozen in liquid nitrogen and stored at -80℃. Crude lipids were extracted using a modified Folch method (Folch et al., 1957). Tissues were homogenized in ice-cold chloroform:methanol (2:1) and incubated at room temperature for 2 hours. Following addition of 0.4 volumes 0.9% sodium chloride, each sample was centrifuged at 2000g at 4 ℃ for 5 min. The organic phase was removed, dehydrated with flow nitrogen overnight, and resuspended in a butanol:Triton-X:methanol solution (0.6/0.267/0.133). Cholesterol content was determined using an enzymatic colorimetric cholesterol assay (Infinity™ Cholesterol Liquid Stable Reagent, Thermo Fisher Scientific, Waltham, Massachusetts #TR13421) and normalized to wet tissue weight.

#### Immunolabeling of fixed frozen sections

Mice were anesthetized and perfused transcardially with 1X PBS followed by 4% paraformaldehyde (PFA) in PBS, brains were post-fixed for 2hrs in the same fixative followed by 48hrs in 30% sucrose/PBS. Half brains were divided through midline and embedded in OCT and frozen at -80C°. 12 μm brain coronal sections were collected using a cryostat (Cryostar NX50, Thermo Fisher Scientific, Waltham, Massachusetts) from the rostral cortical areas (+2.2 mm/+1.2 mm from Bregma) and from the hippocampus (-1.2mm/-2.2 mm from Bregma). ^23^ Sections adhered to positively charged slides, were permeabilized and blocked using 1% Triton-X with 10% normal goat serum in PBS at room temperature for 1 hr and incubated with primary antibodies to GFAP (1:1000 Abcam, Cambridge, Massachusetts USA, #Ab4674), Iba-1 (1:1000 Wako, Richmond, Virginia USA, #019–19741), MBP (1:500 Abcam, Cambridge, Massachusetts USA, #Ab40390), MOG (1:500, Sigma-Aldrich, St. Louis, Missouri USA, #MAB5680), NeuN (1:800 Cell Signaling Technology, Danvers, Massachusetts USA, #94403S). synaptophysin (1:1000, Abcam, Cambridge, Massachusetts USA, #ab32127), 4-HNE (1:500, Abcam, Cambridge, Massachusetts USA, #ab46545), TREM2 (1:200, R&D system, Minneapolis, Minnesota USA, #AF1729), C3 (1:10, Abcam, Cambridge, Massachusetts USA, #ab11862), PLP (1:2, AA3 Rat hybridoma), S100β (1:200, Abcam, Cambridge, Massachusetts USA, #ab52642), 6E10 (1:500, BioLegend, San Diego, California USA, #803001) overnight at 4°C. For ASPA (1:750, GeneTex, Irvine, California USA, #GTX110699) staining, sections were delipidated for 10min in 100% Ethanol, rinsed three times with PBS, treated with L.A.B solution for 10 min and rinsed twice with PBS prior to the incubation with blocking solution (10% Normal Goat Serum, 2% BSA, 0.1% Triton X-100). Anti-mouse Alexa Fluor (AF) 488, anti-rabbit AF 568 and 594, anti-sheep AF 647, anti-rat AF 488 and 647 conjugated secondary Ab (1:300 Jackson ImmunoResearch, West Grove, Pennsylvania USA # 115-545-062, # 112-575-143, #111-585-003, #713-605-147, #112-545-003, #112-605-003), and anti-chicken AF 633 (1:300 Thermo Fisher Scientific, Waltham, Massachusetts #a21103) were reacted for 1 hr at RT. Sections were washed and mounted with ProLong Gold Antifade Mounting Medium (Invitrogen, #P36930) and DAPI (SouthernBiotech, Birmingham, Alabama USA, #0100-20) and cured for 24 hrs.

#### Imaging and Data Analysis

Imaging was performed using the Nikon A1R confocal microscope and the Olympus F4000 confocal microscope. A preview image was taken at 2X to select the areas to be imaged at 20 and 40X. All imaging was performed in the same area of the cortex and CA1 in coronal cryosections. We imaged the cortical layer (L)1 to L6 in the somatosensory region of the rostral cortex and the mPFC ^23^. We imaged the CA1 subregion spanning stratum oriens (SO), pyramidalis (SP), stratum radiatum (SR) and lacunaris molecularis (SLM) including the alveus and extending medially to the cingulum and laterally 650μm to include most of CA1 subregion ^23^. We captured one FOV per section imaged at 20X with Z stacks for 10-25 steps with 1 μ step for cellular and membrane staining except for synaptophysin and TREM2 staining which was imaged with 40X objective with 1.5 software zoom with for Olympus and at 60X for Nikon with Z stacks for 10-25 steps with a 1μ step. The Maximum Intensity Projection (MaxIP, NIS Element, Nikon) or Projection Z (cellSENS FV, Evident) was created from the individual Z stack images. 2D image quantification was performed in Fiji using integrated density for myelin membrane markers and for GFAP cellular profiles and manually counting positive cells in cortex or CA1 subregion. Quantification of myelin protein markers, MBP and MOG was performed in a preset ROI by the IntDen feature in Fiji. For CA1, we used an ROI of 400 μm length and 250μm in width across the entire CA1 subregion including alveus, SO, SP, SR and SLM layer. For sensory cortex layer L2-3 and L4-6, we used an ROI of 300 μm length and 250μm of width across the layers. The data were normalized to the MBP, MOG density in 2 months-old CA1 and cortex.

PLP and ASPA were imaged using the Keyence BZ-X810 microscope,1 FOV was imaged at 20x in the somatosensory cortex. Tiled images were taken with an autofocus set for 5 z-slices over 29.9um per image and brightness and black balance of each channel kept the same per section, then aligned per channel and stitched together using the BZ-X800 Imaging software. Images were analysed using Fiji with PLP intensity calculated using 5 equal sized ROIs extending over layers 2 and 3 of the cortex and ASPA was automatically counted in the same ROIs using thresholding and the Binary Watershed process. Both were normalized to total ROI area.

For plaque burden quantification, we included 6E10 stained areas greater than 100 µm^2^ and used the following parameters to quantify: number as the counts of 6E10 stained accumulations per FOV, the area occupied by plaques (%) and the integrated density by using 3D Object Counter v2.0.1 plug-in in Fiji.

The Imaris (v10.0.1, Oxford Instruments, Abingdon, UK) with Surface tracking function for volume analysis and Spots function was used for the analysis of TREM2+ microglia in 6 months images (Fig. 2 and SFig. 2) and synaptophysin density (Fig. 2). For TREM2+ microglia, DAPI, Iba1 and 6E10 were rendered by Surface function and adjusted with the quality filter to best match the confocal image. 6E10 staining <1000 µm^3^ in volume was excluded from analysis during surface creation. TREM2 was rendered using the spot feature with an estimated radius of 4 µm and TREM2+ /Iba1+cells were identified by filtering with the distance of the spot to the nucleus set at 3 µm. The TREM2+NPAM was obtained by subtracting the TREM2+ PAM defined by direct contact to the 6E10 staining from the total TREM2+ microglia. To quantify TREM2+ microglia at the plaque niche, we defined the plaque niche as a circle of 50 µm radius around the 6E10 staining. Three such plaque niches were examined manually to count TREM2+/Iba1+ cells (Fig. 2).

Synaptophysin was rendered as spots at 2 µm in diameter and filtered using the quality filter to best align with fluorescent signal. The number of synaptophysin spots was measured at the plaque niche (defined above) and in a comparable size area free of 6E10 staining. The ratio of synaptophysin spots at plaque to those in the plaque free area was used to calculate the percent loss.

For 4-HNE+ neurons, DAPI and NeuN were rendered by Surface function and adjusted with the quality filter to best match the confocal image. 4-HNE stained membrane structures were rendered using the spot feature with an estimated radius of 2 µm and 4-HNE+/NeuN+ cells were identified by filtering with the distance of the spot to the nucleus set at 1 µm. For C3+ astrocytes, DAPI, S100β were rendered by Surface function and adjusted with the quality filter to best match the confocal image. C3 was rendered using the spot feature with an estimated radius of 4 µm and C3+/S100β+ cells were identified by filtering with the distance of the spot to the nucleus set at 3 µm. A similar workflow was established for C3+/GFAP+.

#### Protein Isolation and Western Blot Analysis

Mice were deeply anesthetized, perfused with cold PBS and rapidly decapitated. Brains were removed and prefrontal cortices and hippocampi were carefully dissected out on ice ^85^. All tissues were rapidly frozen in liquid nitrogen and stored at -80C°. Frozen tissue was placed into a 1.5mL tube containing homogenization buffer (25mM Tris-HCL pH 7.4, 1mM EDTA, 150mM NaCl, 1% SDS, Protease (Complete Mini, EDTA free, Roche, #11836170001) and Phosphatase inhibitors (PosSTOP, Roche, #04906837001)), and disrupted with a motorized pestle, then further sonicated briefly. Samples were centrifuged at 12,000 rpm for 20 minutes at 4°C and supernatant was collected. Extracted proteins concentrations were determined using a Pierce BCA Protein Assay Kit (Thermo Fisher, Waltham, Massachusetts) and stored at 80°C. Proteins were separated in 4-12% gradient Bis-Tris gels (NuPage) by SDS-PAGE, transferred to Immobilon FL membranes and reacted with primary Ab PLP (AA3 Rat Mab), MBP (Mouse Mab, Biolegend, # 808402) and αtubulin (Mouse Mab, Sigma-Aldrich, #T5168) and labeled with IR Dye 680RD/800CW secondary’s (Li-Cor Biosciences). Membranes were imaged using Odyssey Image Studio software and quantified in FIJI. Loading accuracy was determined using α tubulin as internal control.

### 2.8 Hypoxia Assay. Pimonidazole

To assess areas of hypoxia, we have used the Hypoxyprobe (2-nitroimidazole hypoxia marker known as pimonidazole) kit (HP2-100, Hypoxyprobe, Inc., Burlington, Massachusetts) following the manufacturer’s instructions. Pimonidazole (60 mg/kg) was injected i.p. and four hours after injection mice were anesthetized with 2% isoflurane and perfused intracardially with PBS and 4% PFA. Brains were post-fixed for 24 hours in 4% PFA, equilibrated in 30% sucrose and embedded in OCT. 12 m cryosections were collected from the hippocampus (-1.2/-2.2 from Bregma) and cortex (+2.2/+1.2 from Bregma) and stained with the FITC-conjugated Ab supplied in the kit which detects conjugates of pimonidazole with protein SH groups in hypoxic cells. 20X confocal images were acquired in a Nikon A1R microscope by z stacking. We used two quantification approaches: in one, the fluorescence per area was measured by integrated density with Fiji, in the second, we counted the number of fluorescently labelled cells identified by DAPI in the CA1 subregion and normalized by the volume.

#### Statistics and Reproducibility

For behavioral testing, the experimenter was blinded to the identity of the subjects. The number of subjects tested was 7-17 per genotype and the ratio of M:F was 60:40 at 6 months, 50:50 at 12 months and 40:60 at 16 months. Data were generated by the EthoVision XT 9.0 (Noldus Information Technology, Wageningen, Netherlands) for NSR and Y maze and plotted, as described ^25,80^. For cholesterol quantification, the hippocampus and cortex (+2.2/+1.2 Bregma) from one hemisphere were collected from 3-8 subjects per genotype. Whenever possible for quantitative imaging, the experimenters were blinded to the identity of the samples. For all imaging datasets, we analysed two to three sections per mouse over a set of four mice per genotype. Each technical replicate was analysed separately and averaged to obtain the mean per biological replicate. The biological replicates were then pooled for final genotype analysis. Sample sizes and number of mice were based on means and SEM generated in our laboratory and our published work ^17,86,87^. We did not use statistical methods to predetermine the sample size. After data acquisition and processing, the data were plotted in GraphPad by T-HT or ALS. All data are expressed as mean±SEM. One-way ANOVA with Tukey’s post hoc test or unpaired t-test when only two group comparison was used. All statistical analyses were performed in GraphPad Prism Version 10.4.1. by T-HT or ALS.

#### Cell suspension preparation and scRNA-seq library generation

Mice were deeply anesthetized and perfused intracardially with ice cold PBS, brains were removed, and hippocampi were quickly dissected on ice. We collected hippocampi from 2 brains/sample. The cell suspensions were prepared using the Adult Brain Dissociation Kit (Miltenyi Biotec 130-107-677) according to the manufacturer’s instructions with the following modifications that increased cell yield. The tissues were chopped into small pieces on ice prior to adding to the Enzyme 1 mix, after enzymatic and mechanical dissociation, the mixtures were triturated with P1000 5-10 times to dissociate undigested pieces of tissue, red blood cell removal step was omitted since brains were thoroughly perfused with PBS. Cell counts and viability were measured with AO/PI staining in an automated cell count system (Cellometer Cell Viability Counter, Nexcelom). Cells were immediately fixed using 10X FLEX reagents and 10x Genomics protocol, counted post-fixation and stored at -80 C according to 10X Genomics protocol. 4 biological replicates per genotype (M:F ratio was 40:60), each replicate containing hippocampi from two brains, were submitted to the Single Cell Transcriptomics Core at the University of Pittsburgh. Each sample contained ∼300,000 cells and the viability was >80%. Libraries were prepared using the Chromium Fixed RNA Profiling Reagents Kit for Multiplexed Samples (10x Genomics) according to the manufacturer’s protocol. Libraries sequencing performed by the Health Sciences Sequencing Core at the University of Pittsburgh generated 1.8 billion reads which covers more than 10,000 reads per cell.

#### SCT data processing

Raw FASTQ sequencing files were demultiplexed and aligned to the Chromium Mouse Transcriptome Probe Set v1.0.1 (mm10-2020A) from 10x Genomics using the Cell Ranger multi pipeline (version 8.0.1). The resulting filtered feature-barcode matrices were imported and merged using the Scanpy Python toolkit for downstream analysis. Initial quality control (QC) involved filtering out low-quality cells based on gene detection thresholds. Specifically, cells with fewer than 350 detected genes (n_genes_by_counts) were excluded, with this threshold determined from the distribution of gene counts across all cells. In addition, cells exhibiting high mitochondrial gene expression—defined as exceeding five median absolute deviations (MADs) above the median—were removed to eliminate stressed or dying cells. To further improve data quality, potential doublets were identified and excluded using the scDblFinder function from the scDblFinder package (version 1.20.2), with the parameter dbr.sd set to 1. All cells flagged as doublets were removed from subsequent analyses.

#### SCT integration and clustering

For dataset integration, we employed the scVI pipeline using an iterative approach to enhance cluster resolution and eliminate technical artifacts. In the first iteration, the top 2000 highly variable genes were selected using the sc.pp.highly_variable_genes() function with the flavor parameter set to “seurat_v3”. The scVI model was then trained using sample as a categorical covariate and n_genes_by_counts and pct_counts_mt as a continuous covariate. Clustering was performed using the Leiden algorithm with a resolution of 1.0, followed by a second round of subclustering using the same parameters to increase granularity. After each clustering round, we calculated cluster-level mean QC metrics including the number of genes per cell, total UMI counts, mitochondrial gene percentage, and scDblFinder-assigned doublet scores. Clusters exhibiting high mitochondrial content—defined as exceeding the 99th percentile across all clusters—or mean doublet scores above 0.1 were flagged and removed. This iterative refinement was performed twice to exclude low-quality or artifactual clusters. However, biologically relevant intermediate states, such as those representing oligodendrocyte precursor cell (OPC) to oligodendrocyte (Oligo) lineages, which were sometimes flagged as doublets, were retained. To better resolve microglial and mature oligodendrocyte (MOL) subpopulations, we used the subclusters derived from the second iteration as the starting point. Clusters lacking distinct marker gene expression profiles were systematically merged using the *merge_clusters* function from the transcriptomic clustering package, which implements the split-and-merge strategy.^88^. To identify marker genes specific to each cluster, we used the FindAllMarkers ^89^function implemented in Seurat V5 with the parameters: only.pos = TRUE, min.pct = 0.25, logfc.threshold = 0.25, and max.cells.per.ident = 500.

#### Cell type annotation

Cell type annotation was conducted using the MapMyCells platform (https://knowledge.brain-map.org/mapmycells/process/), which enables reference-based classification of single-cell transcriptomic data. As a reference taxonomy, we employed the 10x Genomics Whole Mouse Brain dataset. Annotation was performed using the Hierarchical Mapping algorithm, which assigns cell identities across multiple hierarchical levels of brain cell types. For major cell type (MCT) classification, final identities were determined by integrating annotations across these hierarchical levels provided by MapMyCells. At the high-resolution cell type (HCT) level, we further refined annotations by incorporating curated subclusters of microglia and mature oligodendrocytes (MOLs). This refinement was guided by a split-and-merge clustering strategy, expression profiles of canonical marker genes, and distinct transcriptional signatures associated with individual clusters.

#### Ambient RNA removal

Potential ambient RNA contamination was removed using decontX^90^, and the corrected expression matrix was subsequently used for downstream differential gene expression analyses and marker gene identification.

#### Differential Gene Expression and Functional Gene Ontology Enrichment Analysis

Differential gene expression (DGE) analysis was performed using a pseudobulk-based approach implemented in muscat (v.1.20.0) ^91^, employing the default edgeR method. Raw counts were aggregated at the cell type level to generate pseudobulks across celltypes. DGE analysis was conducted by comparing genotypes (experimental groups), while controlling for age and sex as covariates. Genes identified as significantly differentially expressed were subsequently used for functional enrichment analysis. Gene Ontology (GO) enrichment was carried out using the Metascape ^92^online tool (https://metascape.org/gp/index.html#/main/step1).

#### Cell type composition analysis

Cell fractions were calculated for each sample. Cell types with fewer than 20 cells per experimental group were excluded from the analysis. A generalized linear model (GLM) was fitted using the glm function in R, with age and sex as covariates. P-values were adjusted using the Benjamini-Hochberg (BH) method via the p.adjust function, treating cell type as a factor to account for multiple testing across cell types.

#### Figure Production

Representative images were rendered in either Imaris or ImageJ, brightness and contrast were altered across the image for better viewing.

## Data Availability

Raw and processed data will be made available in GEO.

## Statement of authorship

T-H.T., F.C. planned the experiments, T-H.T. performed and analysed behavioural and biochemical experiments, T-H.T., XT. T., P. R. and M. R. performed cryosectioning, and immunohistochemistry. T-H. T., A.L.S. and M. R. performed confocal analysis and quantitative image analysis. XT. T., P. R., V. F. generated the new line and managed mouse colonies, T-H.T. performed all statistical analysis, prepared figures. S.B. processed raw single-cell RNA-seq data, performed computational analyses, prepared figures related to the scRNA-seq results, A. S. assisted with computational data analysis. F. C. designed all studies with input from H.M. for the SCT, from J.B.G. for myelin analysis and from T.D.K. for overall conceptualization. F.C. and T.D.K. obtained funding for the study. F.C. wrote the manuscript with input from T-H. T, S.B and H.M. T.D.K., J.B.G., H.M. T-H. T. and S.B. provided input on data interpretation. T.S. and T.C.S. provided the App^NL-G-F^ line. All authors reviewed and edited the manuscript.

## Acknowledgments

This work was supported by NIH (R03 AG072218, R01 NS129632, F.C. and T.D.K.), (R01NS105691, NIH R01NS115707, NSF CBET CAREER 1943906, T.D.K.) (1R01MH 098742, 1R01MH126773, J.B.G.). Part of the confocal imaging was performed in the University of Pittsburgh Dietrich School Microscopy and Imaging Suite (RRID:SCR_022084). The single cell libraries construction was performed at the Single Cell Core Facility (RRID:SCR_025110). Services and instruments used in this project were graciously supported, in part, by the University of Pittsburgh the Office of the Senior Vice Chancellor for Health Sciences. Sequencing was performed on a NextSeq 2000 P4 100 flow cell at the Health Sciences Sequencing Core at UPMC Children’s Hospital of Pittsburgh Rangos Research Center (RRID:SCR_023116). Services and instruments used in this project were graciously supported, in part, by the University of Pittsburgh, the Office of the Senior Vice Chancellor for Health Sciences, the Department of Pediatrics, the Institute for Precision Medicine, and the Richard K Mellon Foundation for Pediatric Research.

