## Supplementary figures and images for "*Fus* depleted oligodendrocytes reduce neuronal damage and attenuate AD progression in the App^NL-G-F^ mouse"

### SFigure 1-anxiety.jpg

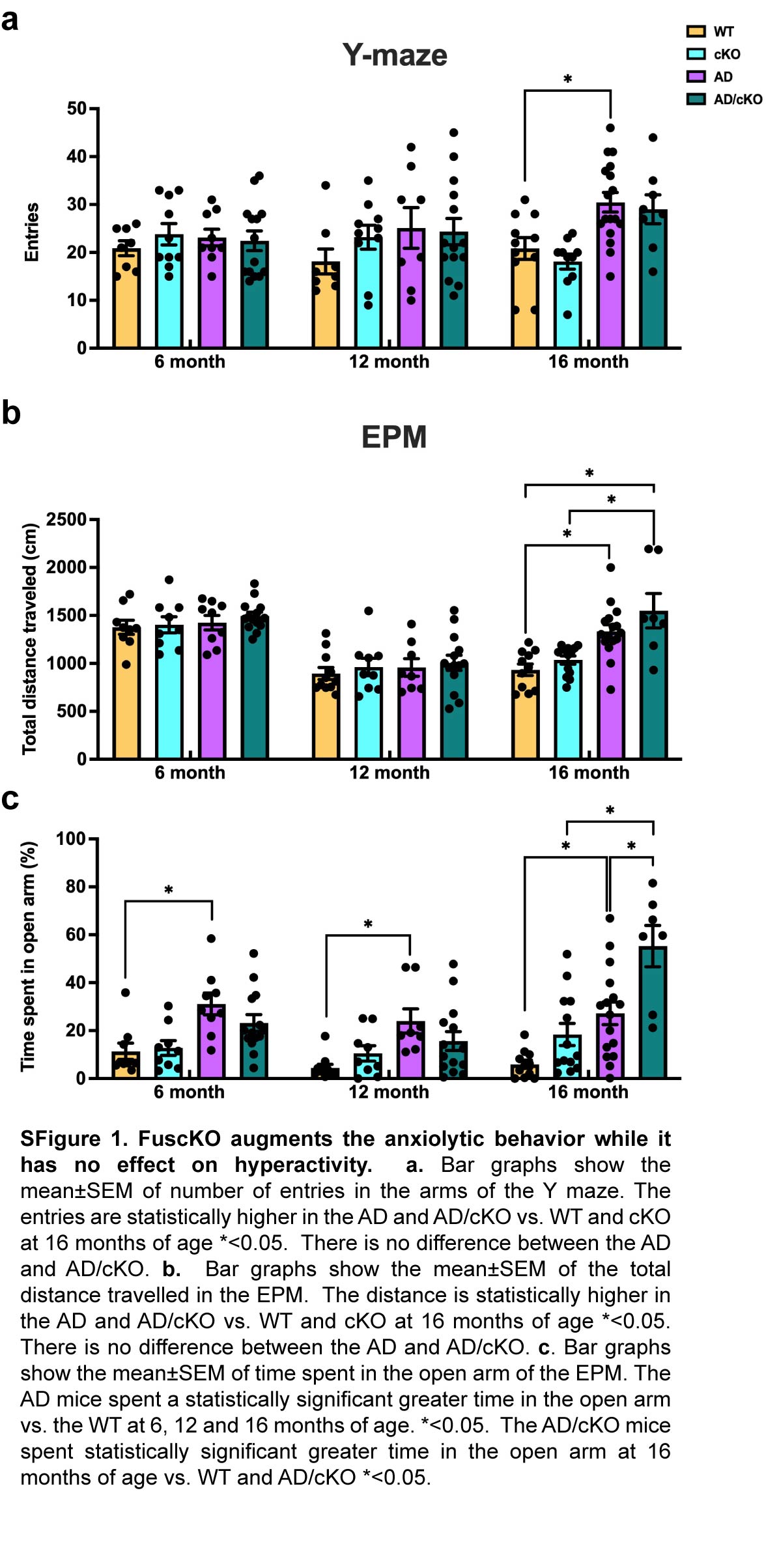

### SFigure 2 -TREM2 PAM.jpg

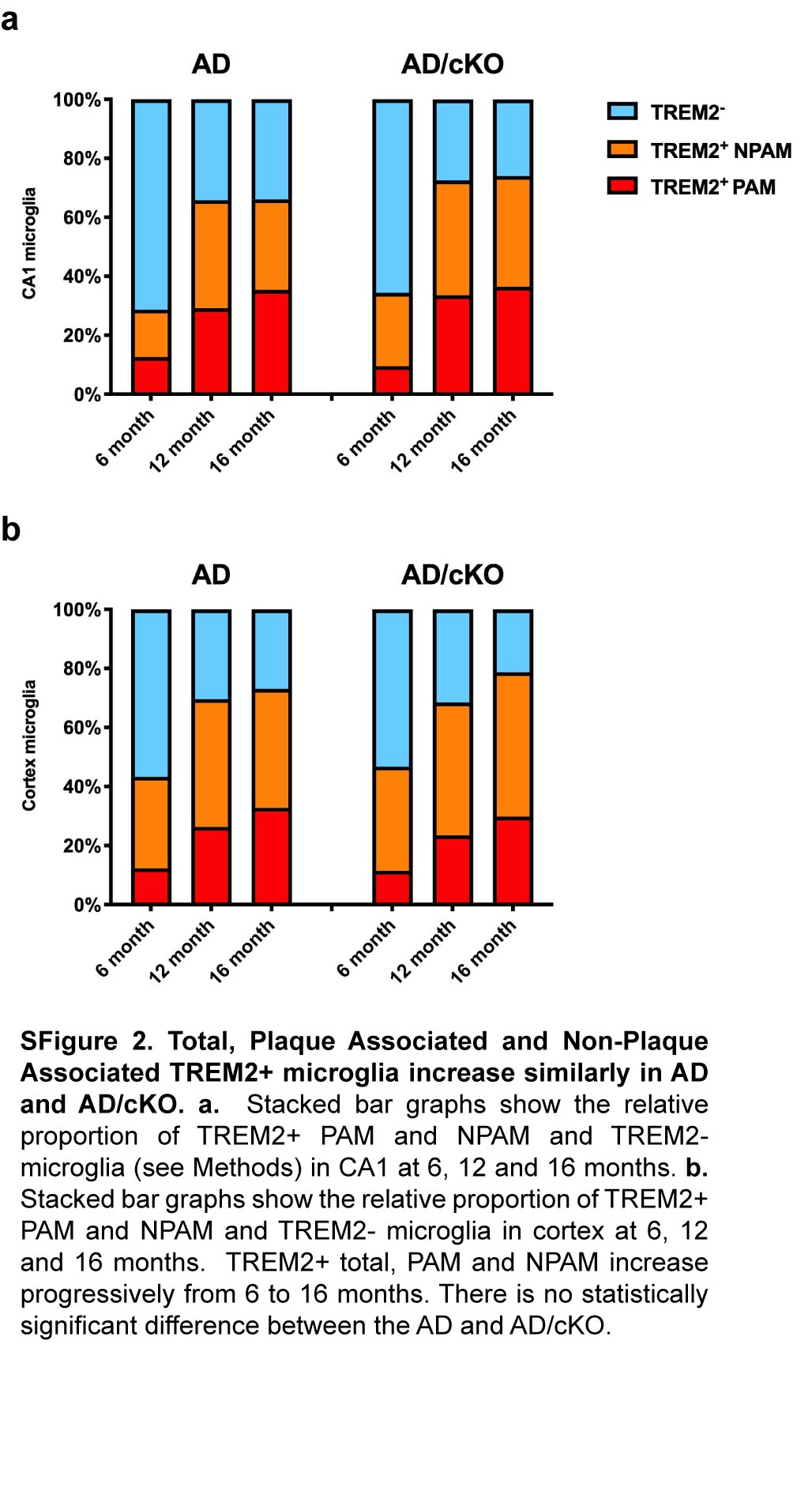

### SFigure 3_L4-6 myelin and WB.jpg

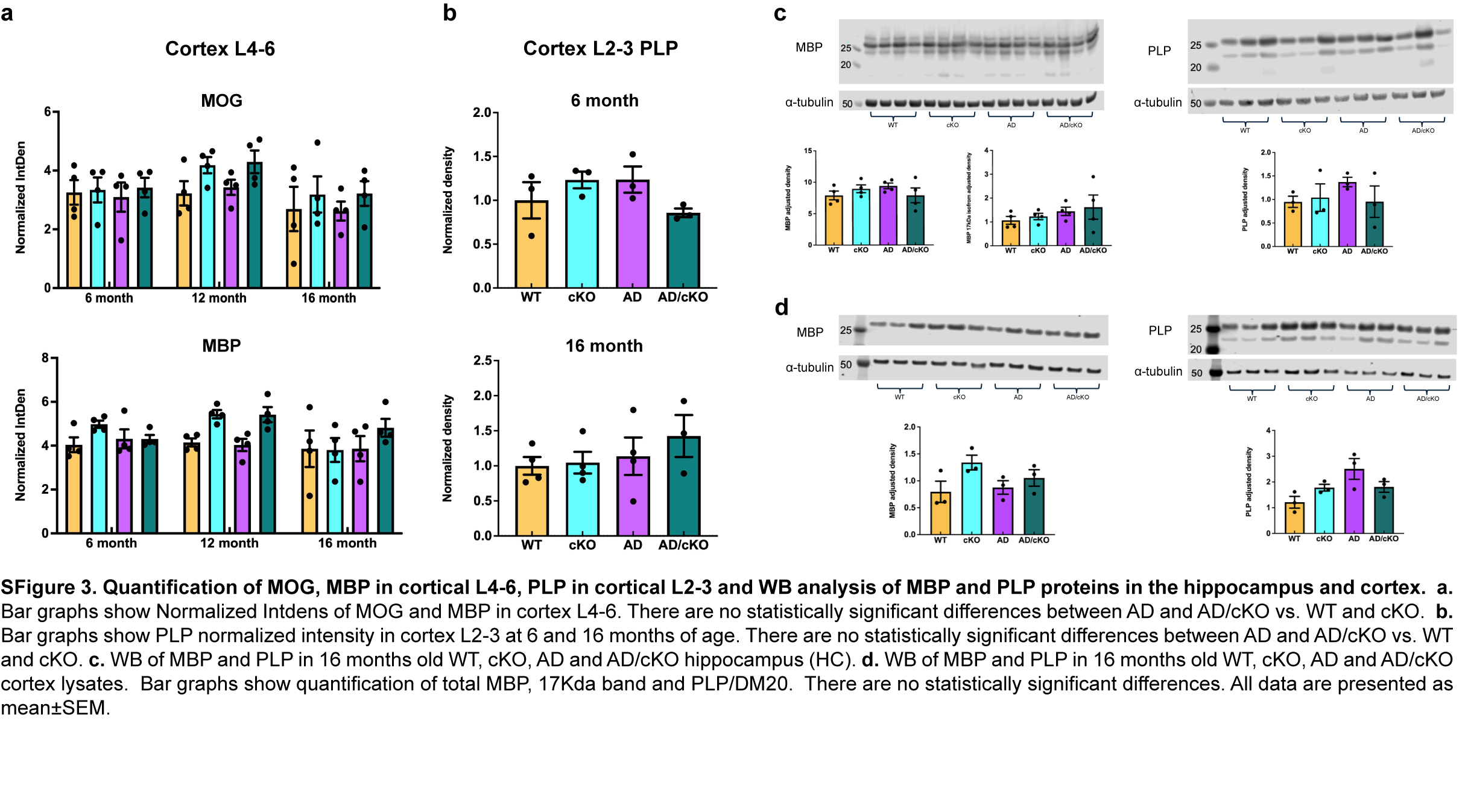

### SFigure 4-mPFC data.jpg

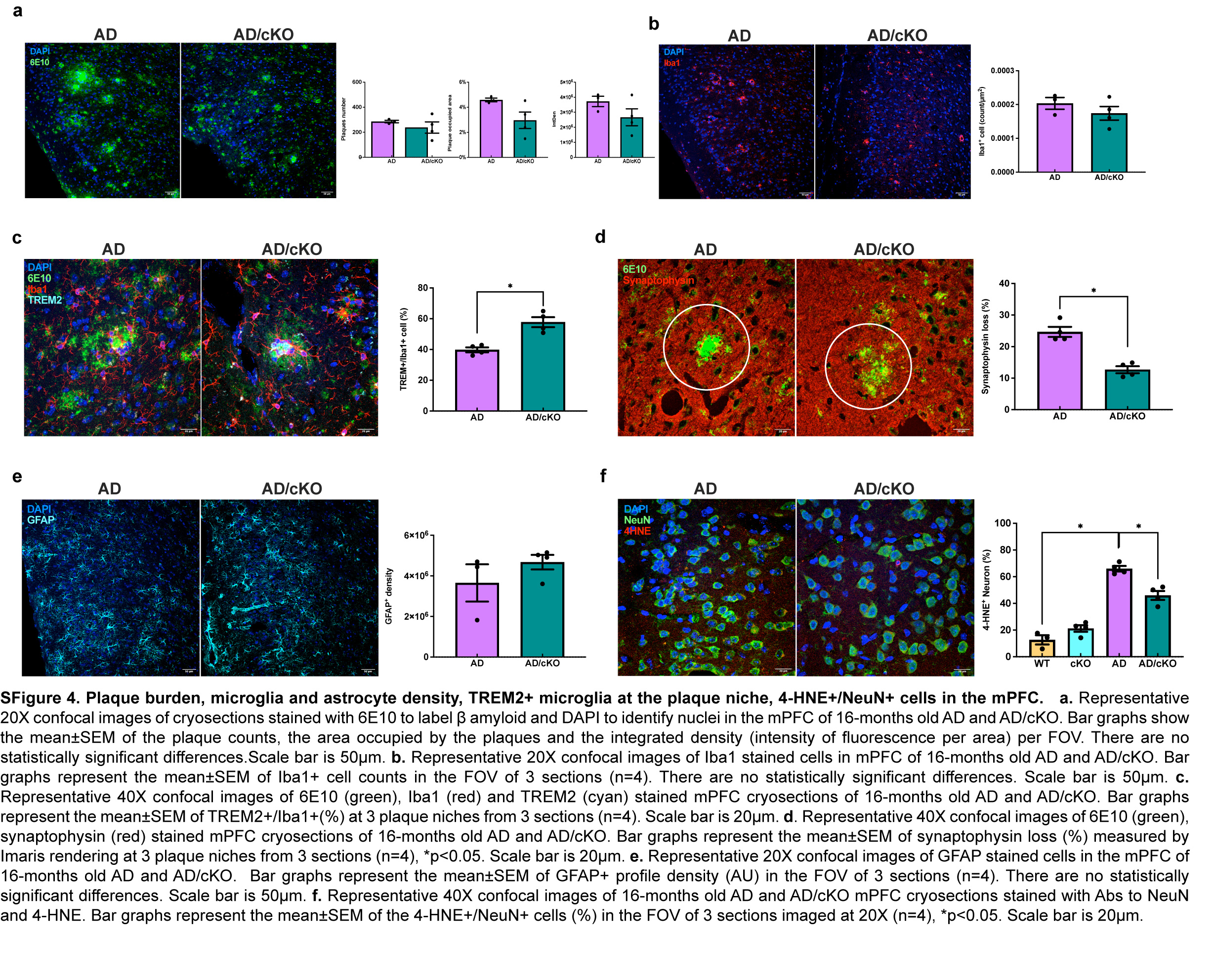

### SFigure 5-SCT2.jpg

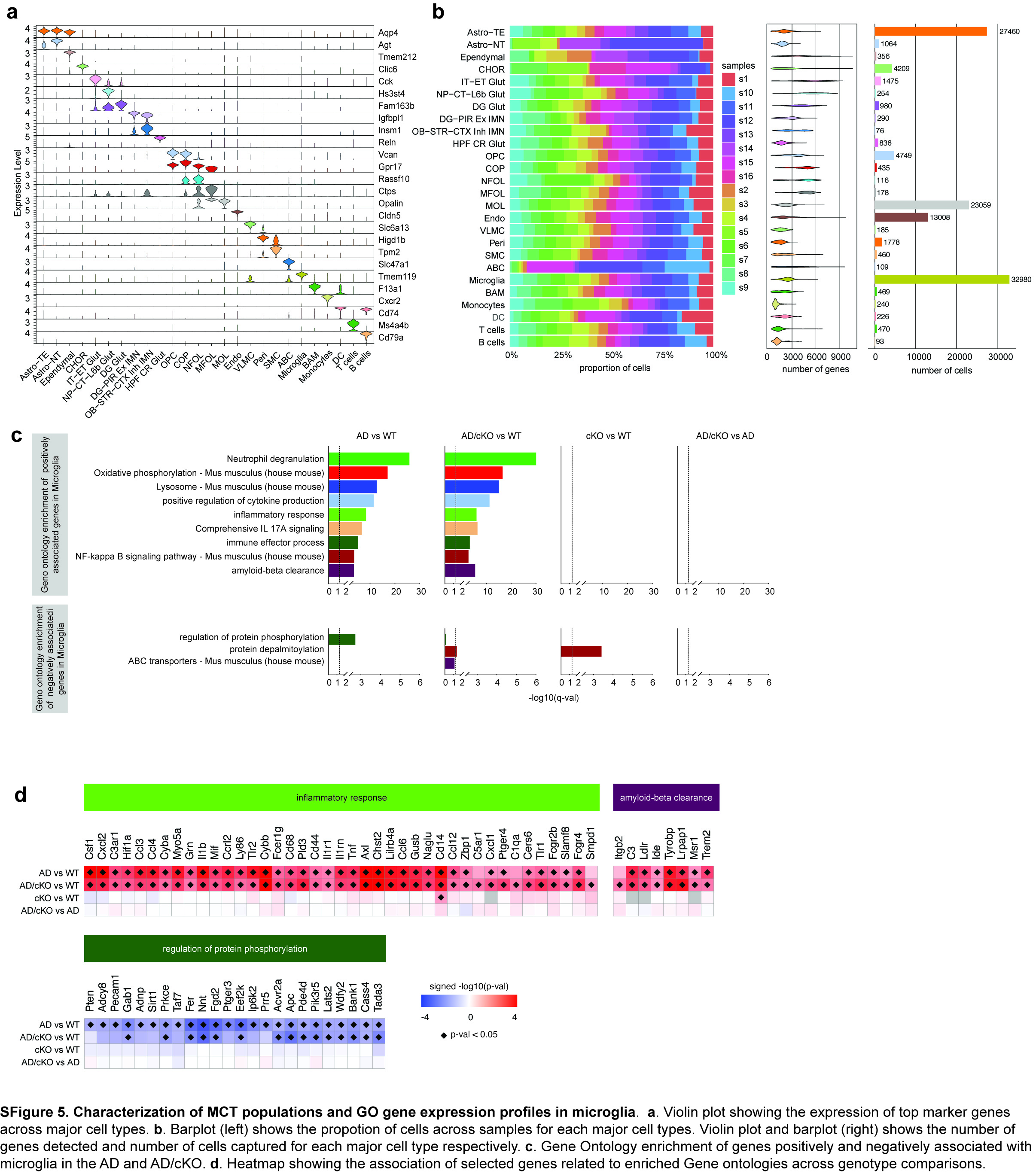

### SFigure 6-SCT3.jpg

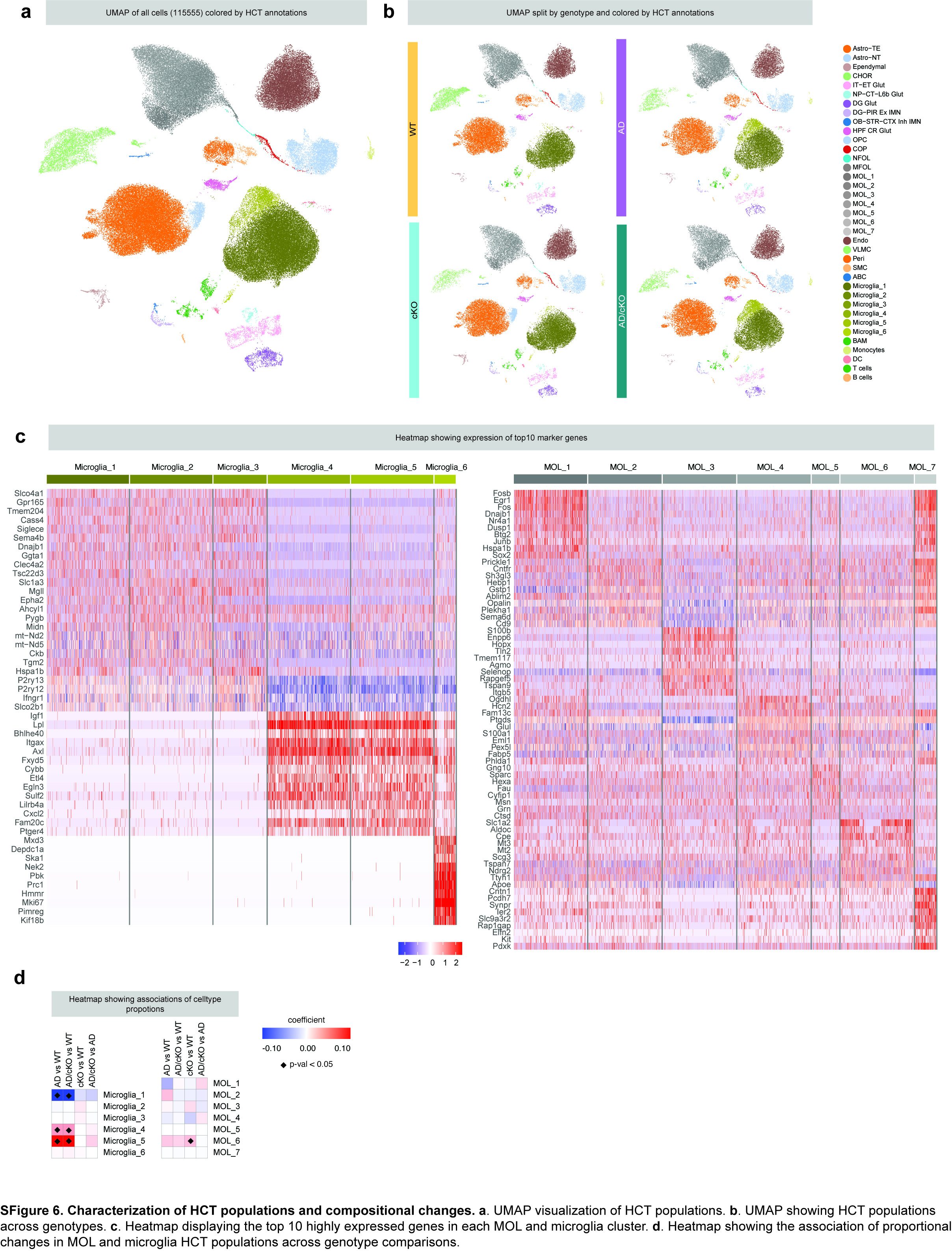
